# Population imaging of neural activity in awake behaving mice in multiple brain regions

**DOI:** 10.1101/616094

**Authors:** Kiryl D. Piatkevich, Seth Bensussen, Hua-an Tseng, Sanaya N. Shroff, Violeta Gisselle Lopez-Huerta, Demian Park, Erica E. Jung, Or A. Shemesh, Christoph Straub, Howard J. Gritton, Michael F. Romano, Emma Costa, Bernardo L. Sabatini, Zhanyan Fu, Edward S. Boyden, Xue Han

## Abstract

A longstanding goal in neuroscience has been to image membrane voltage, with high temporal precision and sensitivity, in awake behaving mammals. Here, we report a genetically encoded voltage indicator, SomArchon, which exhibits millisecond response times and compatibility with optogenetic control, and which increases the sensitivity, signal-to-noise ratio, and number of neurons observable, by manyfold over previous reagents. SomArchon only requires conventional one-photon microscopy to achieve these high performance characteristics. These improvements enable population analysis of neural activity, both at the subthreshold and spiking levels, in multiple brain regions – cortex, hippocampus, and striatum – of awake behaving mice. Using SomArchon, we detect both positive and negative responses of striatal neurons during movement, highlighting the power of voltage imaging to reveal bidirectional modulation. We also examine how the intracellular subthreshold theta oscillations of hippocampal neurons govern spike output, finding that nearby cells can exhibit highly correlated subthreshold activities, even as they generate highly divergent spiking patterns.

## Introduction

Imaging of the activity of neurons using genetically-encoded fluorescent calcium indicators offers the ability to survey neural activity in densely labeled neural circuits, but lacks the temporal precision and sensitivity of less-scalable techniques such as patch clamp electrode neural recording. Due to the desire to be able to image population neural activity with high temporal precision and sensitivity, much recent interest has focused on the development of genetically-encoded voltage indicators (GEVIs). Near-infrared fluorescent GEVIs derived from rhodopsins offer high temporal fidelity, and are compatible with optogenetics^1–3^, whereas green fluorescent GEVIs derived from voltage sensing domains of phosphatases or opsins are slower and brighter^4–9^. Translating these into the living mouse brain has been challenging, because poor membrane localization, photostability, and sensitivity of previous molecules has resulted in poor signal-to-noise ratio (SNR) *in vivo.* In published work, only Ace2N has been used to optically report voltage dynamics in multiple single cells in a living mouse brain, reporting only a single example of the activities from just two cells in one field of view^9^.

Recently, we developed a robotic directed evolution approach that could perform multidimensional optimization of fluorescent voltage indicators along the axes of photostability, sensitivity, and membrane localization, resulting in the rhodopsin-based voltage sensor Archon1, which performed well in the *C. elegans* nervous system, larval zebrafish brain, and in mouse brain slice^3^. Here, we report a variant we call SomArchon, which exhibits improved SNR sufficient to achieve a key milestone: the ability to report the spiking and subthreshold activity of 4-8 neurons at once, in multiple brain regions of awake behaving mice, using inexpensive and simple conventional one-photon microscopes. Making SomArchon turned out to be fairly straightforward, and builds from prior work on targeting opsins to neural somata^10–13^: we conducted a screen for peptides to localize Archon1 to the soma, causing increased signal and reduced background (by eliminating neuropil signals from indicator-expressing axons and dendrites that corrupt the imaging of somatic voltage transients; **Supplementary Figure 1**). The impact was significant, resulting in improvements in sensitivity, SNR, and the number of neurons imageable by manyfold in each case, over prior published reagents. Taking advantage of this improved SNR, we were able to use simple one-photon fluorescence optics to perform wide-field voltage imaging in multiple cortical (visual and motor cortex) and subcortical (hippocampus and striatum) brain regions in awake, attentive and/or behaving mice. We were able to detect neurons in the striatum whose activities were negatively modulated by movement, previously reported by electrophysiology but not easily detected using modern calcium imaging techniques^14–16^, and highlighting the complexity of how the striatum encodes movement. We were also able to measure the phase of spiking relative to intracellular theta oscillations in hippocampal CA1 neurons, confirming earlier experimental results using *in vivo* patch clamping, and further extending these studies by examining how the spiking output of nearby neurons varied despite highly coherent intracellular subthreshold oscillations.

## Results

To target Archon1 to the cell body, we chose amino acid sequences that were shown in the past to target microbial opsins, popular for optogenetic control of neural activity, to the cell bodies of neurons. We chose six such localization motifs and fused them with Archon1 for further validation (see **Supplementary Table 1** for the design of the fusion constructs and the full amino acid sequences of the selected motifs). The selected motifs were fragments of the N-terminus of the kainate receptor subunit KA2 (*ref*^10^), a 222-amino-acid intracellular loop between transmembrane domains I and II of the NaV1.2 sodium channel^17^, a 27-amino-acid segment within the intracellular loop between transmembrane domains II and III of the Nav1.6 sodium channel^17–19^, a 65-amino-acid segment at the C terminus of the Kv2.1 potassium channel^13^, and the membrane-binding domain of the adaptor protein ankyrinG^20^. Additionally, we tested other ankyrinG domains, such as the spectrin-binding domain (AnkSB-motif), the tail domain (AnkTail-motif), the COOH-terminal domain (AnkCt-motif), and the serine-rich domain (AnkSR-motif), that were shown to be sufficient to restrict GFP localization to cell bodies and axon proximal segments^21^. Screening constructs in primary cultured mouse neurons with wide-field fluorescence microscopy, we found that 4 out of 12 constructs showed fluorescence changes during spontaneous neuronal activity (**Supplementary Table 1**). The Archon1-KGC-EGFP-Kv2.1-motif-ER2 fusion protein exhibited the highest ΔF/F during 100-mV voltage steps and appeared to have good soma localization. We named this molecule SomArchon.

SomArchon was functional in mouse brain slice after *in utero* electroporation into mouse cortex and hippocampus (**Supplementary Figure 2**), and after adeno-associated virus (AAV)-mediated expression in mouse cortex, striatum, and thalamus (**Supplementary Figure 3**), revealing spiking of multiple individual cells at once on a conventional one-photon microscope equipped with a sCMOS camera. Quantification of SomArchon localization in neurons in mouse brain slices revealed that SomArchon fluorescence was negligible >30 μm away from the cell body in neurons of the mouse cortex and striatum, and >45 μm in mouse hippocampus, while still demonstrating excellent membrane localization (quantification, **Supplementary Figure 4**; histology of Archon1 vs. SomArchon imaged under identical conditions, **Supplementary Figure 5, 6**). In short, we were able to target Archon1 to the cell body in multiple regions of the mouse brain (**Figure 1a**).

**Figure 1.**
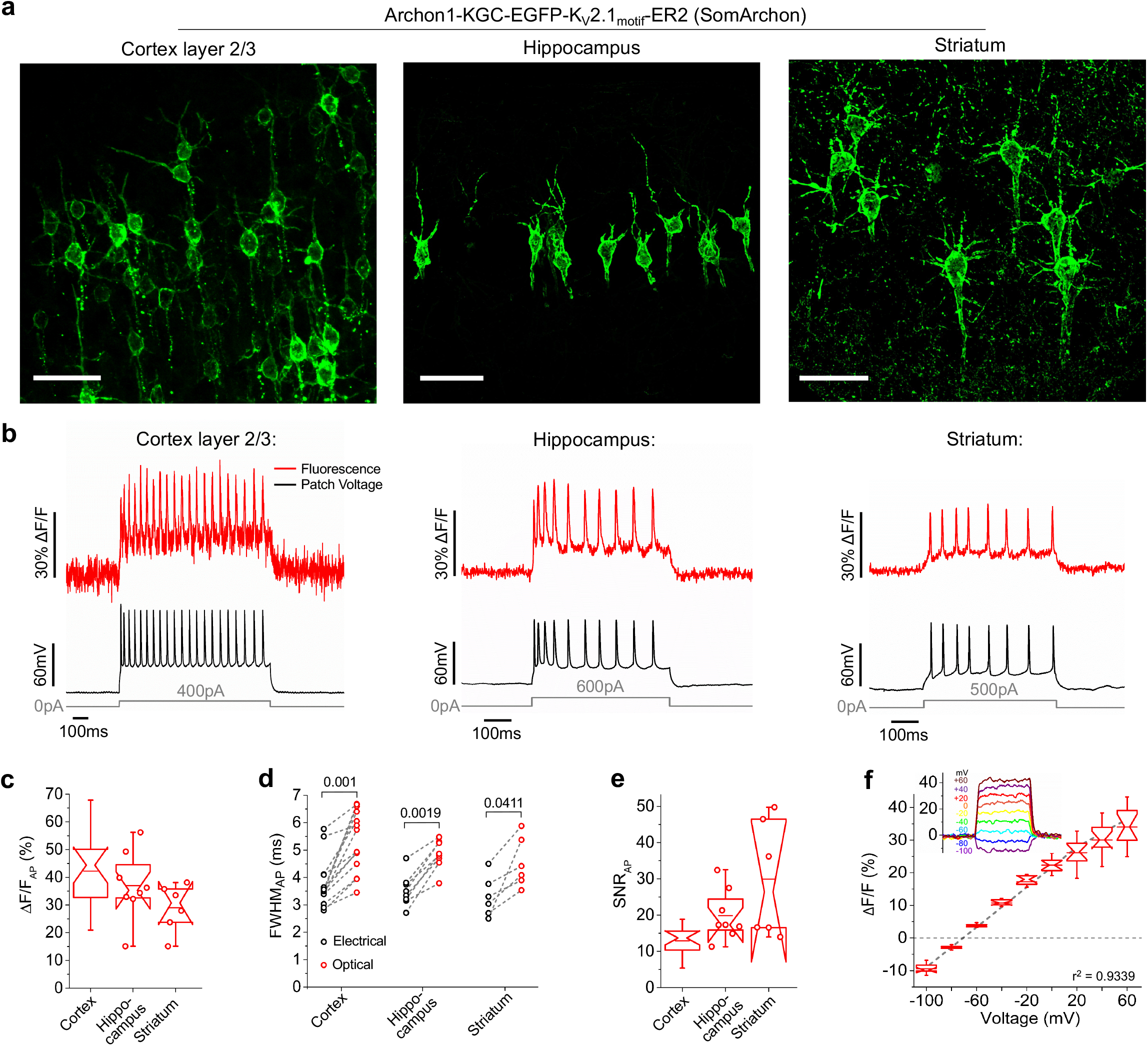
SomArchon enables high fidelity neural voltage imaging in brain slices. (**a**) Representative confocal images of neurons in cortex layer 2/3 (left), hippocampus (middle), and striatum (right) expressing Archon1-KGC-EGFP-Kv2.1_motif_-ER2 (SomArchon for short). Images acquired via EGFP fluorescence using laser excitation at 488 nm and emission 525/50 nm. Scale bars, 50 μm. (**b**) Single-trial optical recording of SomArchon fluorescence responses (red) during current injection through the recording pipette (gray) resulting in action potentials (APs), concurrently recorded via whole-cell patch-clamp in current-clamp mode (black). Imaging conditions throughout the figure: 637-nm laser excitation at 0.8, 1.5, and 1.5 W/mm^2^ for cortex layer 2/3, hippocampus, and striatum, respectively; emission 664 nm longpass; image acquisition rate: ~1 kHz using EMCCD for cortex layer 2/3 and sCMOS for hippocampus and striatum. (**c**) Quantification of ΔF/F per AP across all recordings performed as shown in **b** (n = 18, 8, and 6 neurons from 5, 2, and 2 mice for cortex layer 2/3, hippocampus, and striatum, respectively). Box plots with notches are used throughout this paper (narrow part of notch, median; top and bottom of the notch, 95% confidence interval for the median; top and bottom horizontal lines, 25th and 75th percentiles for the data; whiskers extend 1.5x the interquartile range from the 25th and 75th percentiles; horizontal line, mean; individual data points shown as open circles when n < 9). (**d**) Quantification of electrical and optical waveform full width at half maximum (FWHM; dashed lines connect data points from same neuron) per action potential (AP) across all recordings performed as shown in **b** (n = 14, 8, and 6 neurons from 5, 2, and 2 mice for cortex layer 2/3, hippocampus, and striatum, respectively; *p*-value indicated is from two-sided Wilcoxon rank sum test between electrical and optical waveform FWHM). (**e**) Quantification of signal-to-noise ratio (SNR) per AP across all recordings performed as shown in **b** (n = 18, 8, and 6 neurons from 5, 2, and 2 mice for cortex, hippocampus, and striatum, respectively). (**f**) Population data corresponding to fluorescence changes of SomArchon in response to a series of voltage steps in voltage-clamp mode (optical recordings for a representative neuron shown in inset) recorded in neurons in cortex layer 2/3 (n=12 neurons from 2 mice). Data was normalized so that −70 mV was set to 0 ΔF/F.

We evaluated the performance of SomArchon in multiple regions of the mouse brain via simultaneous patch clamp electrophysiology and one-photon imaging in brain slice (**Figure 1b**). The voltage sensitivity of SomArchon obtained was ΔF/F of 42 ± 12%, 37 ± 11%, and 26 ± 7% (mean ± standard deviation throughout) per action potential for cortex layer 2/3, hippocampus, and striatum, respectively (**Figure 1c**; imaging conditions described in the caption; n = 18, 8, 6 neurons from 5, 2, 2 mice for cortex layer 2/3, hippocampus, and striatum, respectively). This represents an improvement over the parent, Archon1, of ~2-fold in sensitivity, likely because of decreased background neuropil fluorescence (compared to the mouse cortex brain slice data of Piatkevich *et. al.* 2018^3^). SomArchon fluorescence followed high-speed changes in voltage during single action potentials, with fluorescence waveforms being only slightly broader than electrically-recorded action potential waveforms (**Figure 1d**, n = 14, 8, and 6 neurons from 5, 2, and 2 mice for cortex layer 2/3, hippocampus, and striatum, respectively; **Supplementary Table 2**). During action potentials, SomArchon fluorescence changes exhibited SNRs (defined as the maximum voltage change observed during a recording divided by the standard deviation of the baseline) of 13 ± 4, 20 ± 6, and 27 ± 13 per action potential, in the cortex, hippocampus, and striatum, respectively (**Figure 1e**; imaging conditions described in the caption; n = 18, 8, 6 neurons from 5, 2, 2 mice for cortex layer 2/3, hippocampus, and striatum, respectively). This is about the same as the parent Archon1 (compared to mouse cortex brain slice data in Piatkevich et. al. 2018^3^). In addition, the dependence of SomArchon fluorescence on voltage was linear (r^2^ = 0.9339) throughout the physiological range of membrane potentials from −100 mV to +60 mV (**Figure 1f**; n = 12 neurons from 2 mice), with sufficient sensitivity to report subthreshold changes. SomArchon expression did not alter membrane resistance, membrane capacitance, or the resting potential of neurons in the cortex, hippocampus, and striatum in comparison to the control neurons recorded in acute brain slices (**Supplementary Figure 7, Supplementary Table 2**).

In work being considered in parallel to this manuscript, the molecule SomaQuasAr3 (**Supplementary Figure 8**), a soma-localized version of an improved version of the QuasAr2 sensor, was developed^22^. We did a side-by-side comparison with SomArchon under identical conditions, using the blue-light driven channelrhodopsin CoChR to elicit action potentials in cortex layer 2/3. Under the same imaging conditions we used for cortical neurons above, i.e. 0.8 W/mm^2^ of excitation intensity at 1 kHz acquisition rate, we could not detect any functional activity of SomaQuasAr3. To increase the SNR for SomaQuasAr3 and enable spike detection, we raised the light power to 1.5 W/mm^2^ and decreased the acquisition rate to 500-600 Hz. Under these imaging conditions, the ΔF/F per action potential for SomaQuasAr3 was 14 ± 7%, about ¼ of the 54 ± 30% observed for SomArchon. The SNR per action potential for SomaQuasAr3 was 6 ± 2, about 1/6 of the 37 ± 16 for SomArchon (n = 9 and 14 cells from 2 mice each for SomaQuasAr3 and SomArchon, respectively; **Supplementary Figure 8**). Thus, SomArchon achieves a ~5x benefit in sensitivity and SNR over molecules being considered in parallel to this one.

One can compare a new fluorescent indicator to older ones in terms of quantitative metrics as well as qualitative ones, e.g. what new kinds of science are enabled. We compared ΔF/F and SNR of SomArchon to those values for all of the recently published genetically-encoded voltage indicators that were imaged under wide-field microscopy in acute mouse brain slice, namely Ace2N-mNeon^9^, ASAP1^5^, and MacQ-mCitrine^7^ For comparison, ΔF/F per action potential for SomArchon was 4.7-, 7-, and 17-fold higher than that of green fluorescent voltage sensors Ace2N-mNeon, ASAP1, and MacQ-mCitrine, respectively. Although SNR is hard to compare across studies because it depends on camera acquisition rate and excitation light power, the SNR per action potential for MacQ-mCitrine at a light power of 30 mW/mm^2^ and a 440 Hz acquisition rate was 6; for ASAP1 at a light power of 8-50 mW/mm^2^ and a 400 Hz acquisition rate, 7.5 ± 2.5; for SomArchon at 1 kHz (a higher frame rate) and 0.8 W/mm^2^, the SNR (see above) was ~2× better (Ace2N-mNeon SNR in slice was not reported)^3,5,7^. QuasAr2 in hippocampal brain slices showed a ΔF/F of 15% and an SNR of 8.5 ± 2.6 per action potential under the high light power of 8 W/mm^2^ (*ref*^23^), about half that achieved by SomArchon under 5x lower light power (**Figure 1c, e**). Thus, SomArchon represents a manyfold improvement in terms of sensitivity and SNR over previously published state-of-the-art molecules.

### SomArchon enables *in vivo* voltage imaging with single cell, single spike precision using conventional wide-field microscopy

Having established the improved performance of SomArchon over prior molecules in terms of sensitivity and SNR, we next sought to quantify whether this was reflected in improved ability to observe neurons *in vivo*. We targeted multiple brain regions with AAV containing SomArchon under the synapsin promoter, including the motor cortex, visual cortex, striatum, and hippocampus. Mice were head-fixed and awake while under a simple, conventional wide-field one-photon microscope (**Figure 2a**). Using the green fluorescence of SomArchon to identify virally-transfected cells using a low magnification (10x) objective lens (**Figure 2b**), we then performed voltage imaging of near-infrared fluorescence of SomArchon using a 20x objective lens at ~1.6 W/mm^2^ of excitation light or a 40x objective lens at ~4 W/mm^2^ of excitation light at 637 nm. SomArchon baseline fluorescence levels were sufficient to resolve individual cells in the near-infrared voltage channel (**Figure 2c, e, g, i**), with single cell bodies resolvable at depths up to 180 μm below the brain surface, and with most falling at depths of 50-150 μm (for cortex and hippocampus) and 20-150 μm (for striatum; see **Supplementary Figure 9** for histology of mice used for *in vivo* imaging).

**Figure 2.**
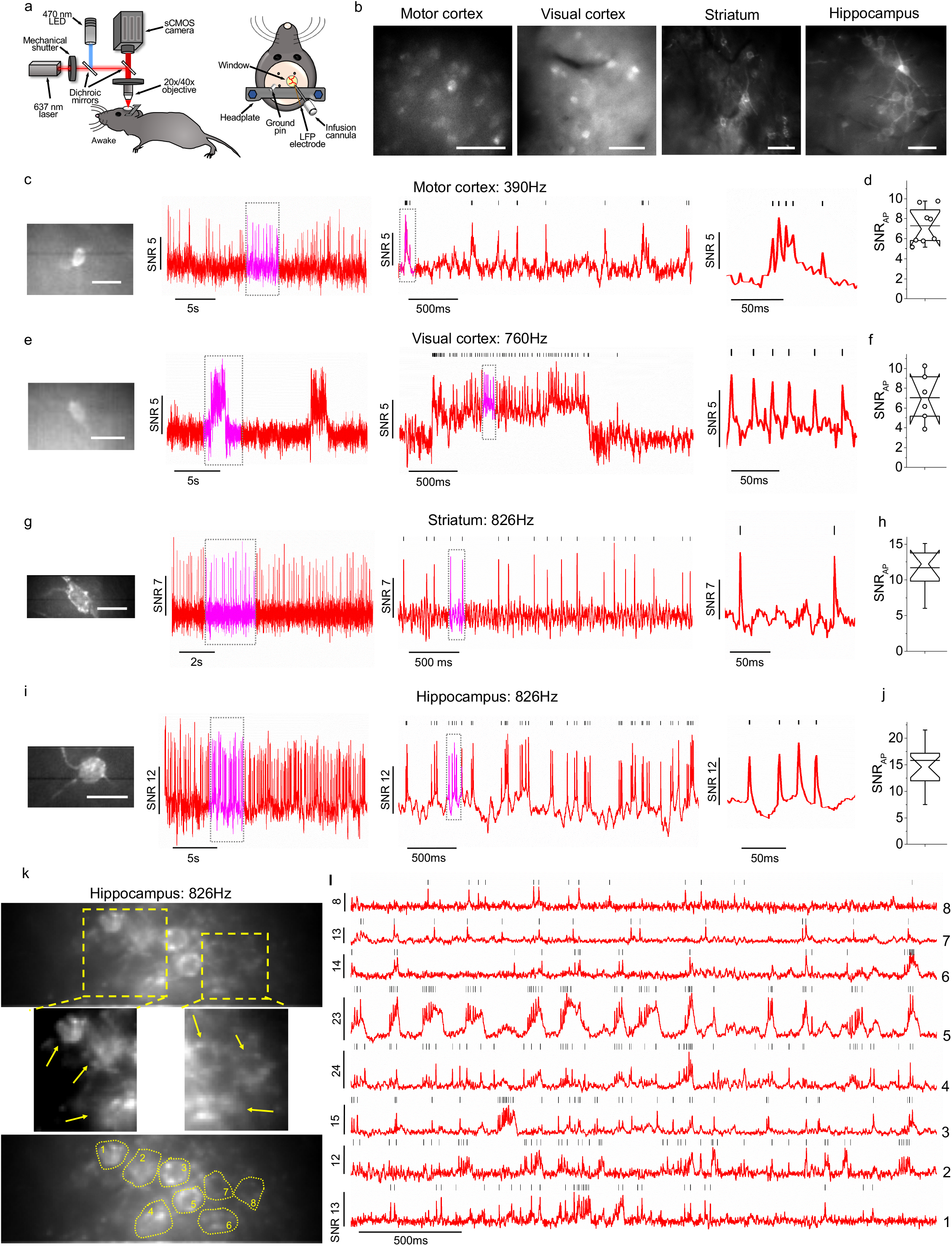
SomArchon-mediated voltage imaging with single cell resolution in multiple brain regions of awake mice, using a simple wide-field imaging setup. (**a**) Schematic representation of the experimental setup: awake mice were head-fixed and positioned under a conventional wide-field microscope that allowed imaging of SomArchon fluorescence in the green and near-infrared channels (*left*); example diagram of surgical window implant (*right*); for striatum and hippocampus, imaging windows were coupled with a viral infusion cannula and a LFP recording electrode). (**b**) SomArchon labeled neurons visualized via EGFP fluorescence in motor cortex, visual cortex, striatum, and hippocampus (ƛ_ex_=470/25 nm LED, ƛ_em_=525/50 nm). Scale bars, 50 μm. (**c, e, g, i**) Voltage imaging in (**c**) motor cortex, (**e**) visual cortex, (**g**) striatum, and (**i**) hippocampus: SomArchon fluorescence image of the cell *in vivo (left)* and optical voltage trace acquired from this cell *(right;* dashed boxes indicate time intervals shown at successively expanded time scales; vertical bars indicate peaks of action potentials identified by spike-sorting algorithm); ƛ_ex_=637 nm laser, ƛ_em_=664LP, excitation intensity 1.5 W/mm^2^ for visual and motor cortex, 4 W/mm^2^ for striatum and hippocampus). Scale bars, 25 μm. (**d, f, h, j**) Quantification of signal-to-noise ratio per action potential for (**d**) motor cortex, (**f**) visual cortex, (**h**) striatum, and (**j**) hippocampus (n = 8, 6, 10, and 17 cells from 3, 2, 3, and 4 mice, respectively). (**k, l**) Population voltage imaging in hippocampus: (**k**) SomArchon fluorescence images of cells *in vivo;* summed fluorescence intensity over ~250ms recording (top), with ROIs overlayed *(bottom)*, zoomed-in views of two regions boxed in *top* to highlight individual cells (*middle*), and (**I**) optical voltage trace acquired from these cells (trace numbers correspond to cells numbers in **k**); ƛ_ex_=637 nm laser, ƛ_em_=664LP, excitation intensity 1.5 W/mm^2^.

Optical voltage recordings of awake, head-fixed mice with an acquisition rate of 390-910 Hz using an sCMOS camera consistently detected individual spikes, as well as bursts of spikes, in single cells across all four brain regions (**Figure 2c, e, g, i**, and **Supplementary Video 1**). Since ΔF/F was less informative across imaging sessions due to the large variation in background fluorescence across imaged fields of view, we quantified the SNR per action potential in the motor cortex, visual cortex, striatum, and hippocampus as 7 ± 2 at ~1.6 W/mm^2^ excitation power (n = 8 neurons from 3 mice), 7 ± 2 at ~1.6 W/mm^2^ excitation power (n = 6 neurons from 2 mice), 12 ± 3 at ~4 W/mm^2^ excitation power (n = 10 neurons from 3 mice), and 16 ± 7 at ~4 W/mm^2^ power (n = 17 neurons from 4 mice), respectively (**Figure 2d, f, h, j**). Unfortunately, no other paper reports SNR values per action potential in the living mouse brain, so we cannot directly compare our molecule to others (and thus the most accurate comparison we can conduct is in the context of brain slice, as noted earlier). In the hippocampus, we were able to resolve proximal dendrites and detect voltage fluctuations in these projections that differed from those in the soma (**Supplementary Figure 10**), which may enable the analysis of dendritic integration and other intracellular computational processes *in vivo*.

Due to the high performance and soma-targeted nature of our construct, it was straightforward to image multiple neurons at once, e.g. 8 spiking neurons at once in the hippocampus at labeling densities where it was possible to manually select ROIs while avoiding overlapping signals (**Figure 2k, l**; motor cortex data in **Supplementary Figure 11**). While the brightest cells in multicell recordings such as **Figure 2k** could be easily identified from a single frame, defining ROIs for dimmer cells adjacent to these bright cells was more challenging due to the wide dynamic range (16 bit) of the raw images. To select ROIs for these dimmer cells, we created max projection and standard deviation images of the entire raw video and stretched their LUTs (look up tables) to enhance visibility.

The previous record for number of cells imaged in a single field of view was just ¼ of that, namely 2, with Ace2N-mNeon^9^. In a paper being considered in parallel to ours, SomaQuasAr3 was used to visualize up to 4 cells in a field of view^22^, using a custom-built imaging setup that combines two-photon and single-photon microscopy modalities with patterned excitation illumination in order to target individual cell bodies^22^. Thus, SomArchon helps achieve a manyfold improvement in the number of spiking cells that can be detected in a field of view over other indicators, while still using a simple, conventional wide-field microscopy setup.

### All-optical electrophysiology *in vivo* using SomArchon in combination with a channelrhodopsin

The use of microbial rhodopsins has become widespread in neuroscience as a method for activating and silencing the neuronal activity of genetically targeted cell populations^24^. Some high-performance channelrhodopsins are activated with blue light, and thus, could in principle be spectrally combined with near-infrared fluorescence indicators such as SomArchon for *in vivo* neural activation and voltage imaging. We used a bicistronic expression system to coexpress SomArchon and the high-performance channelrhodopsin CoChR^25^ in the same cell via a 2A cleavage sequence (**Figure 3a**). Illuminating positive cells in brain tissue with 2 ms blue light pulses at 10 and 20 Hz resulted in the reliable evocation of single action potentials per pulse with a jitter less than 1 ms (n = 8 neurons from 2 mice; **Figure 3b-d**). We expressed SomArchon-P2A-CoChR-Kv2.1_motif_ (with Kv2.1 chosen because it was better at soma-targeting of the CoChR lacking GFP, in contrast to our earlier paper on SoCoChR fused to GFP which was better soma-targeted with a fragment of KA2) via AAV injection in the hippocampus of the mouse brain and optogenetically evoked neural activity in awake mice while imaging voltage of the same subset of cells (**Figure 3e**). Blue light stimulation for durations of 100 ms evoked spiking, as imaged in populations of neurons within the same field of view (**Figure 3f**; this effect was reliable across different fields of view, **Figure 3g**). Thus, we have shown the feasibility of using optogenetic stimulation while optically recording voltage dynamics in several cells simultaneously in an awake mouse.

**Figure 3.**
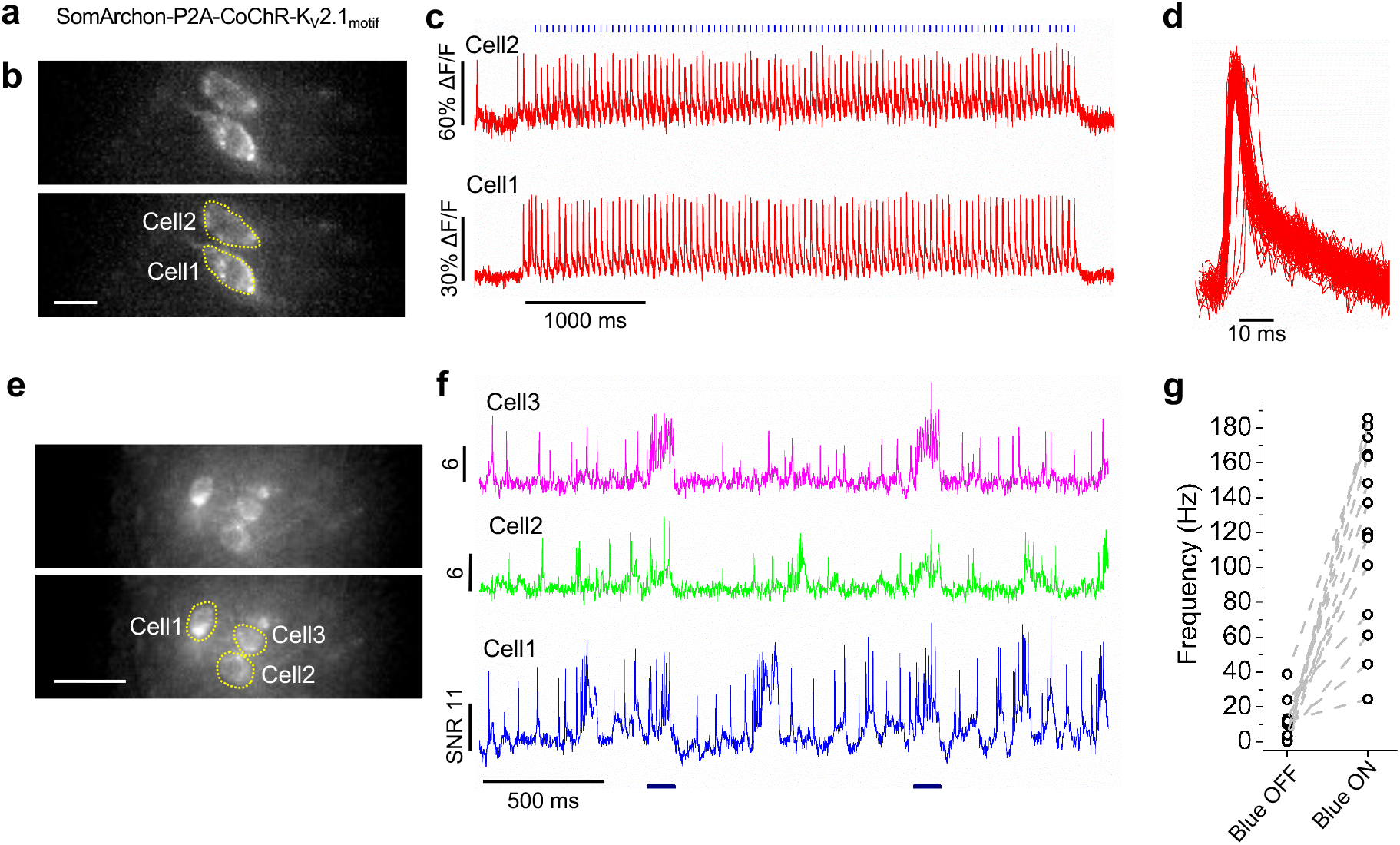
All-optical electrophysiology with CoChR and SomArchon. (**a**) Diagram of the construct used for co-expression of SomArchon and CoChR. (**b**) Fluorescence images of a selected field of view of hippocampal neurons in brain slice expressing SomArchon-P2A-CoChR-Kv2.1 (top) with the identified region of interest (ROI) corresponding to somas shown (bottom); ƛ_ex_=637 nm, ƛ_em_=664LP, exposure time 1.3 ms. Scale bar = 25 μm. (**c**) Representative single-trial optical voltage traces from cells shown in **b** aligned with blue light stimulation (2 ms pulse at 20 Hz). Image acquisition rate: 777 Hz. (**d**) Superposition of single action potentials induced by pulses of blue light for the Cell1 trace shown in **c**. (**e**) Fluorescence images of a selected field of view of hippocampal neurons *in vivo* expressing SomArchon-P2A-CoChR-Kv2.1 *(top)* with the identified ROI corresponding to somas shown *(bottom)*; ƛ_ex_=637 nm, ƛ_em_=664LP, exposure time 1.2 ms. Scale bar = 20 μm. (**f**) Representative single-trial optical voltage traces from cells shown in **e** aligned with blue light stimulation (100 ms pulse). Image acquisition rate: 826 Hz. (**g**) Firing rate changes during blue light off and blue light on conditions in individual neurons (n = 14 cells from 2 mice).

### *In vivo* single cell, single spike optical voltage imaging from striatal neurons during locomotion

Electrophysiology studies have demonstrated that striatal neurons exhibit heterogeneous positive modulations during distinct phases of movement (*i.e.*, encoding discrete aspects of movement initiation, execution, and termination)^26–30^. It has also been reported that some striatal neurons (~5-20%) are negatively modulated during motor behavior^27,29,31^. While electrophysiology recordings largely discard spatial information regarding the relative location of the neurons being observed, recent calcium imaging studies have revealed that spatially clustered neurons encode similar aspects of movement within striatal populations^14–16^. In particular, these studies focus on the increases in activity during movement, perhaps because decreases in activity are hard to observe with calcium imaging due to its slow temporal resolution and therefore, low event rate. Voltage imaging could help resolve these ambiguities by offering the temporal sensitivity and precision of electrophysiology, at the spatial resolution of calcium imaging.

We optically recorded striatal neurons expressing SomArchon while mice ran on a spherical treadmill (**Figure 4a**). When imaged with the 40x objective lens, we could clearly identify individual cell bodies and record voltage dynamics from multiple cells in a single field of view (**Figure 4b, c**), extracting spikes for correlation with movement (**Figure 4d, e**). Individual striatal neurons exhibited highly different firing rates (**Supplementary Figure 12**), with some exhibiting bursting patterns known to occur in striatal fast spiking interneurons^32,33^. We also identified one neuron with a large cell body which exhibited a tonic firing pattern of ~5 Hz (**Supplementary Figure 12**), both properties of cholinergic interneurons^34^.

**Figure 4.**
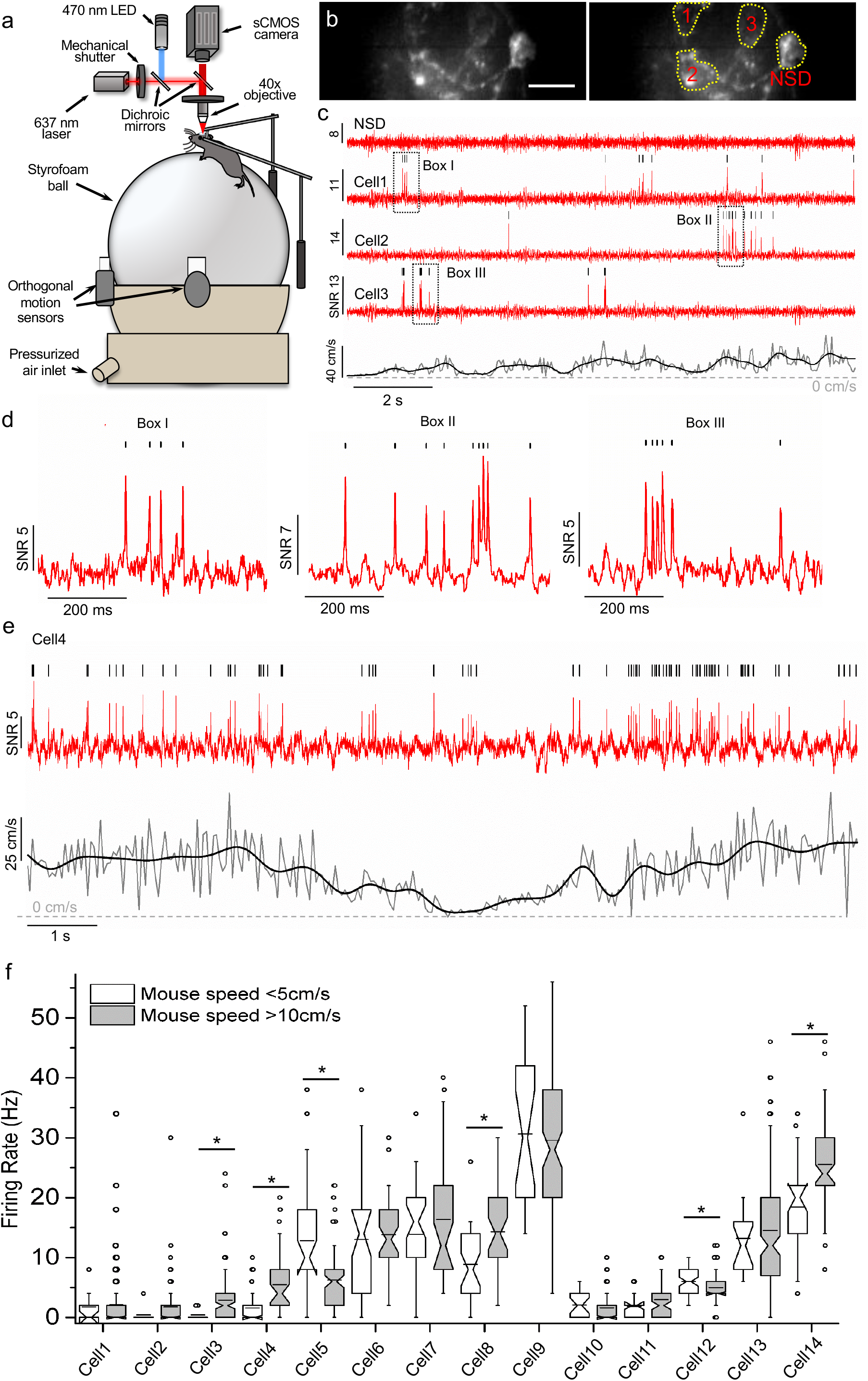
Voltage imaging of striatal neurons during locomotion. (**a**) Schematic representation of the experimental setup: awake mice were head-fixed and positioned under a conventional wide-field microscope identical to that in **Figure 2a**, and positioned on a spherical treadmill. Animal locomotion was monitored using a pair of orthogonally-oriented motion sensors. All imaging was performed with a 40x NA = 0.8 water immersion objective lens. (**b**) SomArchon fluorescence image of striatal cells (*left*), with the identified ROIs corresponding to somas shown (*right*), ƛ_ex_=637 nm, exposure time 1.2 ms. Scale bar, 20 μm. NSD = no spikes detected. (**c**) Representative optical voltage traces acquired from cells shown in **b**, and corresponding mice movement speed (black: low pass filtered at 1.5 Hz; gray: raw data). Image acquisition rate 826 Hz. (**d**) Zoomed-in views of the three periods indicated by black boxes shown in **c**. (**e**) Representative optical voltage trace (red) for a neuron modulated by movement speed; corresponding movement speed (black: low pass filtered at 1.5Hz; gray: raw data). Image acquisition rate 826 Hz. (**f**) Firing rates of individual striatal neurons, during periods with low (open box) versus high (gray) movement speed (open circles: outlier periods). *, *p*<0.05, two-sided Wilcoxon rank sum test.

We segmented each voltage trace into 0.5-second intervals so we could compare firing rate vs. speed for each interval, analyzing periods of high speed movement (speed ≥10 cm/s) and low speed movement (speed ≤5 cm/s) as in previous behavioral studies (see **Supplementary Figure 13** for identification of high and low speed thresholds)^14^. Of 14 neurons recorded, four were positively modulated by movement speed (**Figure 4e, f**; Cells 3, 4, 8, 14; p<0.05 two-sided Wilcoxon rank sum test, **Supplementary Table 2**), and two were negatively modulated by movement speed (**Figure 4f** Cells 5, 12; p<0.05 two-sided Wilcoxon rank sum test). Thus, our data suggest that neurons negatively modulated during movement can be readily detected with voltage imaging.

We found that adjacent neurons did not respond to movement speed in identical ways. For example, in each of the two recordings where three neurons were simultaneously recorded, we found that one of the three neurons was positively modulated by movement speed (p<0.05 twosided Wilcoxon rank sum test, **Supplementary Table 2**), whereas the other two were not modulated at all (**Figure 4b, c, f**; see Cells 1, 2, 3, and Cells 6, 7, 8). Together, these results highlight the potential of SomArchon for recording from spatially clustered striatal neurons during behavior.

### Simultaneous recordings of intracellular membrane potential and LFPs revealed distinct phase relationships between spikes and theta oscillations in hippocampus

We performed wide-field imaging of hippocampal CA1 neurons in awake, head-fixed mice, while simultaneously recording the local field potential (LFP). CA1 neurons exhibited average firing rates ranging from 3.3 to 18.0 Hz (7.92±3.55 Hz, mean± standard deviation, n=16 neurons, from 7 recording sessions from 4 mice), consistent with previous electrophysiology recordings^35,36^. Many prior electrophysiological studies in the hippocampus have established that theta oscillations are prominent in the hippocampus and contribute to many hippocampal-related behaviors^37,38^. Indeed theta oscillations were prominent throughout our recording sessions (**Supplementary Figure 14**). Several theoretical frameworks have been developed to explain how spike-oscillation phase relationships may support hippocampal information processing^39–41^. However, most of these studies compare spikes to extracellularly recorded LFP theta oscillations, leaving open the question of how subthreshold theta oscillations within an individual neuron relates to that neuron’s spiking output.

*In vivo* patch clamp recording has pointed to closer phase-locking of a given neuron’s spikes to its own intracellular theta oscillations, than to the across-neuron averaged LFP theta oscillations^35,42^. We recorded classical electrode-measured LFPs while performing optical voltage imaging in the CA1 region of awake mice, so that we could compare the phase relationships between spikes and intracellularly-measured theta oscillations vs. across-neuron averaged LFP theta oscillations (**Figure 5a**). Some neurons had spikes phase-locked to both intracellular and LFP theta oscillations (**Figure 5a**), whereas others were phase locked only to the intracellular theta oscillations, and not the LFP theta oscillations (**Figure 5b**). We found that 15 out of the 16 observed CA1 neurons showed significant phase locking between spikes and intracellular theta oscillations (**Figure 5c**; blue: p<0.05, cyan: not significant, **χ**^2^ test, comparing to the null hypothesis of a uniform distribution). Spikes were phase locked to the rising phase of intracellular theta oscillations (338±11 degrees, mean± standard deviation, peak of oscillation=0 degree; n=16 neurons). In contrast, only 6 out of 16 neurons showed significant phase locking between spikes and extracellular LFP theta oscillations (**Figure 5c**; red: p<0.05, pink: not significant, **χ**^2^ test, comparing to the null hypothesis of uniform distribution). As a population, neurons exhibited a stronger phase locking relationship with intracellular theta oscillations than with LFP theta oscillations (phase vector for intracellular=0.35±0.17, for LFP=0.12±0.08, mean± standard deviation; **Figure 5d**). These results are consistent with recent *in vivo* whole cell patch clamp recordings in freely moving mice, where spikes were more tightly phase locked to intracellular theta oscillations than LFP theta oscillations^35,42^. Thus, SomArchon can support the analysis of subthreshold intracellular oscillations.

**Figure 5.**
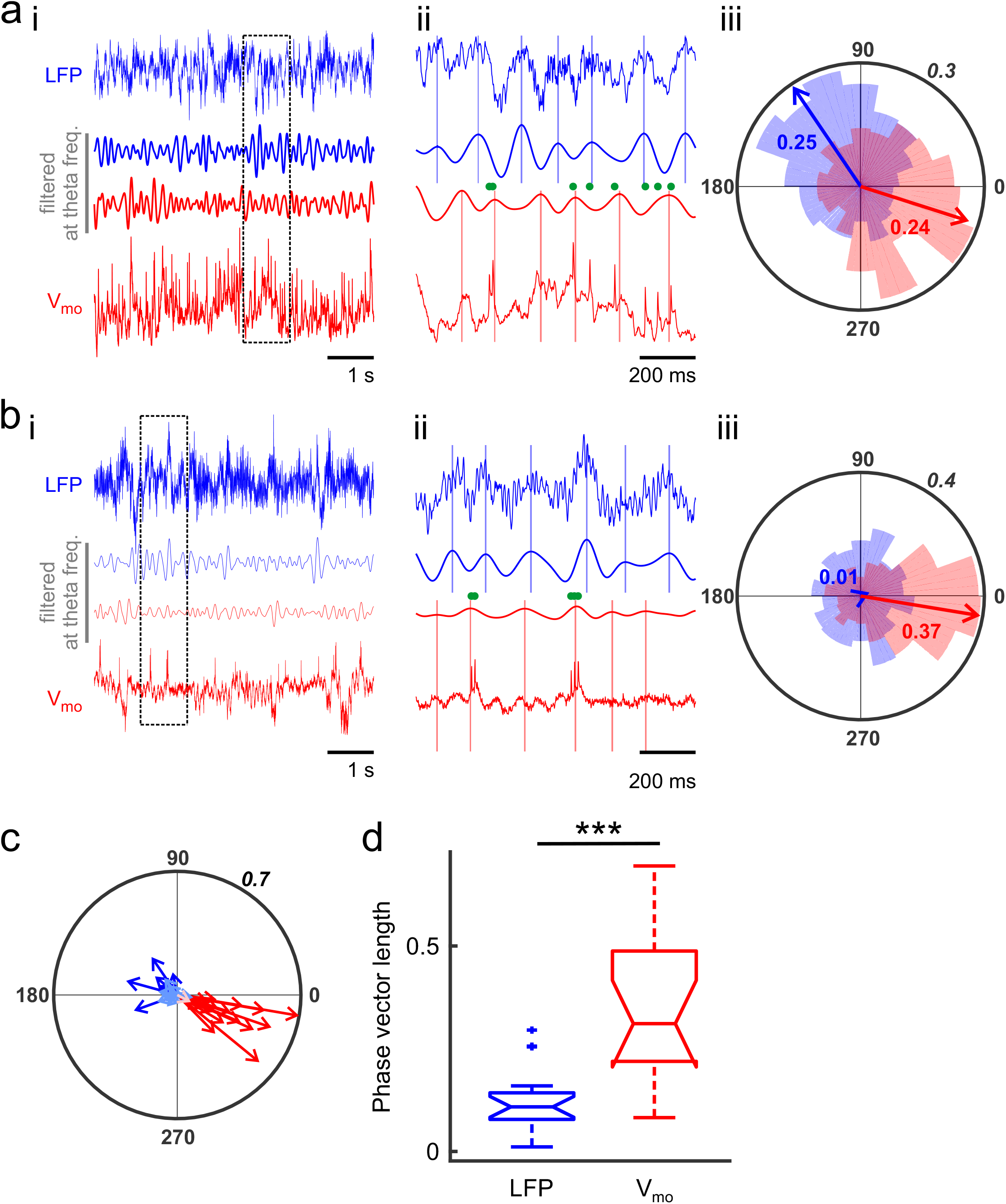
Phase relationships between hippocampal neuron spikes and theta oscillations of optically-recorded membrane voltage (V_mo_) vs. those of local field potential (LFP) recordings. (**a**) An example neuron with spikes phase-locked to theta oscillations of both LFPs and V_mo_. (**i**) Raw LFPs (blue trace, top) and raw V_mo_ (red trace, *bottom)*, as well as theta frequency (4-10Hz) bandpass filtered LFPs (blue, *2^nd^ to the top)* and V_mo_ (red, *2^nd^ from the bottom).* (**ii**) Zoomed-in views of the periods indicated by the black box in **ai**, showing the peak of theta oscillations in LFPs (vertical blue lines) and in V_mo_ (vertical red lines), and the timing of spikes (green dots). (**iii**) Probability distribution of spikes relative to the phases of LFP theta oscillations (blue shaded region) and V_mo_ theta oscillations (red shaded region). Arrows indicate the average phase vector (values next to the phase vectors indicate the vector length). Bin size = 20 degrees. Outer circle indicates probability = 0.3. (**b**) Similar to (**a**), but for an example neuron that is phase locked to V_mo_ theta oscillations, but not to LFP theta oscillations. Outer circle indicates probability = 0.4. (**c**) Population spike phase relationship to LFP theta oscillations (blue and cyan) and V_mo_ theta oscillations (red and pink). Each vector represents the average vector from one neuron. Blue and red: *p*<0.05; cyan and pink: not significant; **χ**^2^ test, comparing to the null hypothesis of uniform distribution. Outer circle indicates probability = 0.7. (**d**) As a population, hippocampal neuron spikes show stronger phase locking to V_mo_ theta oscillations than to LFP theta oscillations (***, *p*<0.001, two-tailed paired Student’s t-test, n=16 neurons in 7 recording sessions from 4 mice). Box plot used in this figure: top and bottom horizontal lines, 25th and 75th percentiles for the data; whiskers extend to the most extreme data points not considered outliers; + are outliers; and horizontal line represents the median.

### Population imaging of spiking and subthreshold oscillations in hippocampal neurons

Having established that SomArchon could report subthreshold intracellular oscillations *in vivo*, we next examined the extent to which they were coordinated across neural populations to regulate spiking. The hippocampus has typically been regarded as having little topographical organization^43,44^, although some evidence has suggested that a degree of spatial clustering exists^45,46^. Recently, based upon multi-neuron automated patch clamp data^47^, we hypothesized that even when nearby neurons in the awake mouse hippocampus exhibit highly coordinated subthreshold activities, they might be capable of generating extremely divergent spiking outputs. Since we were unable to analyze more than one cell in the hippocampus at a time, however, even with automated patch clamp, we were previously unable to test this hypothesis.

We accordingly examined the coherence, both at the spiking level and at the subthreshold intracellular oscillation level, across populations of neurons, both to examine whether nearby neurons exhibited similar subthreshold intracellular oscillations (with a focus on theta frequencies which exhibit strong phase relationship with spikes (**Figure 5c**)), and to examine the extent to which those coordinated activities generated similar spike patterns. In nine hippocampal recording sessions where we recorded multiple neurons, we found that simultaneously recorded pairs of neurons (**Figure 6a**) could exhibit high coherence of subthreshold intracellular oscillations, but low coherence of spiking (**Figure 6b**, green vs. magenta traces). Across the 25 neuron pairs simultaneously imaged, spike-spike coherence was not easily predicted from the subthreshold intracellular oscillation coherence (**Figure 6d**, n = 25 pairs, linear regression, *p* = 0.086, r^2^ = 0. 123). Thus, nearby neurons may receive a common synchronized input, but a small number of selective inputs may have an outsized role in governing spiking output, at least in the awake-at-rest state investigated here.

**Figure 6.**
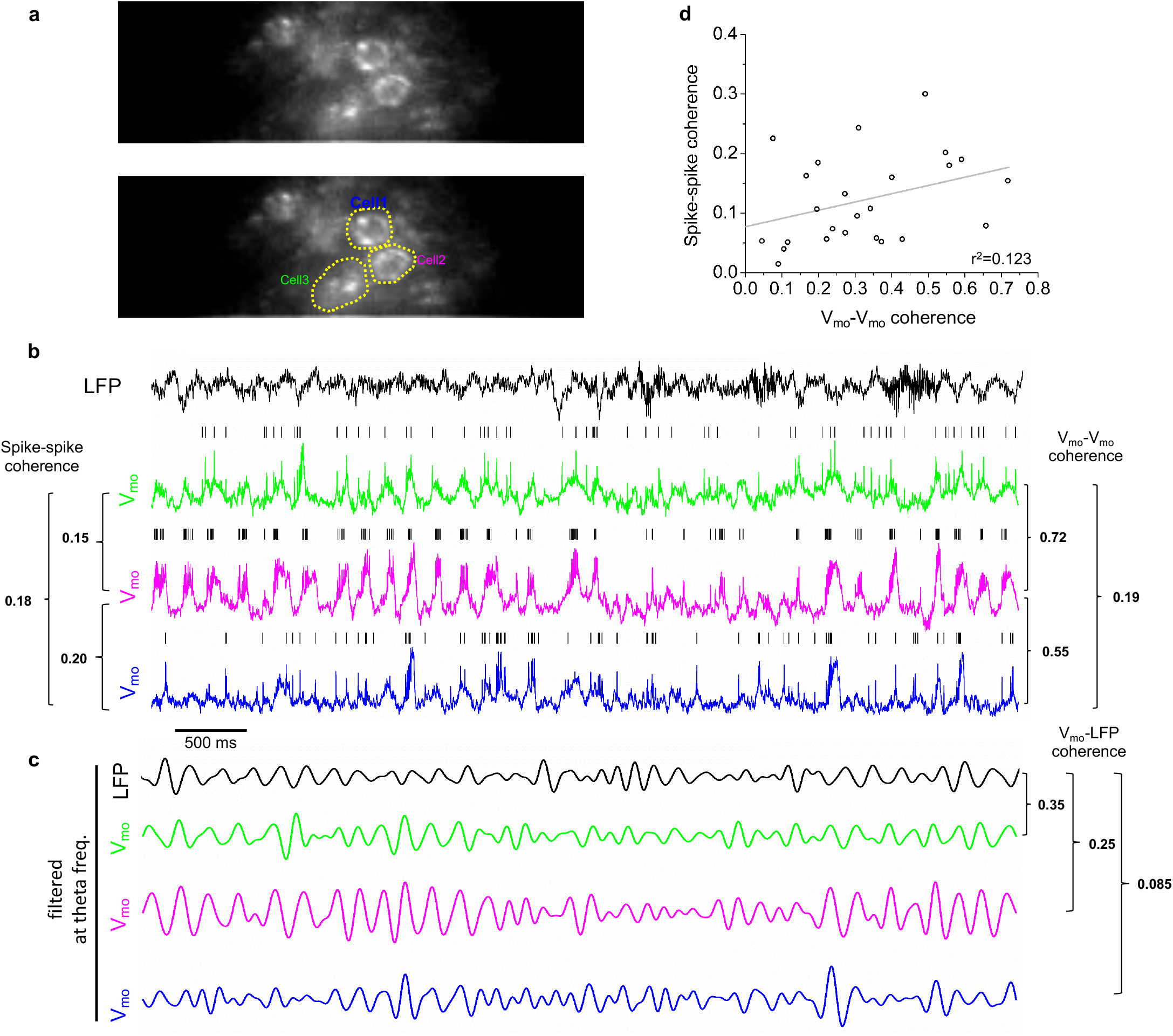
*In vivo* population voltage imaging reveals coupling between intracellular subthreshold oscillations and spiking in hippocampal neurons. (**a**) An example field of view containing three neurons. (**b**) Membrane voltage recorded optically (V_mo_) from neurons identified in **a** (also color coded as in a) and simultaneously recorded local field potentials (LFPs). Black vertical ticks above V_mo_s denote identified spikes. Spike-spike coherence values between neurons are shown at the *left* and V_mo_-V_mo_ coherence values are shown at the *right.* (**c**) Theta frequency-filtered LFPs and V_mo_s for the four traces shown in **b**. V_mo_-LFP coherence values are shown to the *right.* (**d**) Correlation between V_mo_-V_mo_ coherence and spike-spike coherence from all neuron pairs, fitted with a linear regression (n = 25 pairs, *p* = 0.086, r^2^ = 0.123).

The coherence between two nearby neurons’ intracellular subthreshold oscillations was not constant, but fluctuated over time (coefficient of variation, 0.39±0.12, mean±standard deviation) and was significantly different from a stable coherence value (one sample t-test, p<0.001), revealing temporally dynamic input structures. To further explore how intracellular subthreshold oscillations may relate to the across-neuron averaged LFP signals, we examined the coherence between intracellular subthreshold theta oscillations recorded in two neurons, versus the coherence between one neuron and the simultaneously recorded LFP theta oscillations. While simultaneously recorded neurons exhibited a wide range of coherence to LFP oscillations (**Figure 6c**), at a population level, adjacent neurons showed greater intracellular oscillation coherence amongst themselves than with the LFP (V_mo_-V_mo_ coherence=0.32±0.13, V_mo_-LFP coherence=0.10±0.05, mean±standard deviation, two-sided Wilcoxon rank sum test, *p*=0.0008, n = 9 sessions with simultaneous recording of LFPs and multiple neurons; **Supplementary Table 2**, see **Methods**). These results highlight that among the diverse current sources that contribute to the spatiotemporally integrated LFP signals, some may exhibit a greater role in synchronizing post-synaptic responses.

In summary, SomArchon enables the simple, inexpensive observation of neural voltage in populations of individual neurons, in a diversity of brain regions in awake behaving mice. We anticipate that the practicality of SomArchon will enable its rapid deployment into a diversity of contexts in neuroscience. As camera performance improves in years to come, and as further evolution of voltage indicators continues, we anticipate that perhaps dozens to hundreds of neurons will be imageable with simple one-photon optics in the near future.

## Methods

### Molecular cloning

For screening candidates for the soma-localized Archon1 voltage sensor in primary hippocampal neurons, DNA encoding for candidate localization motifs were synthesized *de novo* with mammalian codon optimization and subcloned with Archon1 (GenBank ID MG250280.1) and EGFP genes into the pAAV-CAG vector to obtain the final constructs described in **Supplementary Table 1** (gene synthesis and subcloning were performed by Epoch Life Science Inc.). For *in vivo* expression in the mouse brain via *in utero* electroporation (IUE), the Archon1-KGC-EGFP-Kv2.1_motif_-ER2, QuasAr3-PP-Citrine-Kv2.1_motif_-ER2, and CoChR-mTagBFP2-Kv2.2_motif_-ER2 genes were subcloned into the pCAG-WPRE vector. The QuasAr3-PP-Citrine-Kv2.1-ER2 gene was synthesized de novo (GenScript Biotech Corp.) based on sequences reported in the original preprint^22^. The CoChR-mTagBFP2-Kv2.2_motif_-ER2 gene was assembled by Epoch Life Science Inc. using pAAV-Syn-CoChR-GFP (Addgene plasmid #59070) and pBAD-mTagBFP2 (Addgene plasmid #34632) as the source of the CoChR and mTagBFP2 genes, respectively; the Kv2.2_motif_s were synthesized *de novo* with mammalian codon optimization (Epoch Life Science Inc.). The pAAV-Syn-Archon1-KGC-EGFP-Kv2.1_motif_-P2A-CoChR-Kv2.1_motif_ plasmid was also cloned by Epoch Life Science Inc. Plasmid amplification was performed using Stellar (Clontech Laboratories Inc.) or NEB10-beta (New England BioLabs Inc.) chemically competent *E. coli* cells. Small-scale isolation of plasmid DNA was performed with Mini-Prep kits (Qiagen); large-scale DNA plasmid purification was done with GenElute HP Endotoxin-Free Plasmid Maxiprep Kits (Sigma-Aldrich Corp.).

### Neuronal culture and transfection

All mouse procedures were performed in accordance with the National Institute of Health Guide for Laboratory Animals and approved by the Massachusetts Institute of Technology Institutional Animal Care and Use and Biosafety Committees. For dissociated hippocampal mouse neuron culture preparation, postnatal day 0 or 1 Swiss Webster mice (Taconic Biosciences Inc., Albany, NY) were used as previously described^3^. Briefly, dissected hippocampal tissue was digested with 50 units of papain (Worthington Biochemical Corporation) for 6-8 min at 37 °C, and the digestion was stopped by incubating with ovomucoid trypsin inhibitor (Worthington Biochemical Corporation) for 4 min at 37 °C. Tissue was then gently dissociated with Pasteur pipettes, and dissociated neurons were plated at a density of 20,000-30,000 per glass coverslip coated with Matrigel (BD Biosciences). Neurons were seeded in 100 μL plating medium containing MEM (Life Technologies Corp.), glucose (33 mM, Sigma), transferrin (0.01%, Sigma), Hepes (10mM, Sigma), Glutagro (2 mM, Corning), Insulin (0.13%, Millipore), B27 supplement (2%, Gibco), and heat inactivated FBS (7.5%, Corning). After cell adhesion, additional plating medium was added. AraC (0.002 mM, Sigma) was added when glia density was 50-70% of confluence. Neurons were grown at 37°C and 5% CO2 in a humidified atmosphere.

For *in vitro* screening of candidate soma-localized Archon1 sensors, primary hippocampal neuron cultures were transfected with 500 ng of plasmid DNA per well with a commercial calcium phosphate transfection kit (Life Technologies Corp.) after 4 days *in vitro* (DIV), as previously described. After 30-60 min of DNA-calcium phosphate precipitate incubation with cultured neurons at 37°C, neurons were washed twice with acidic MEM buffer (pH 6.7-6.8) to remove residual calcium phosphate particles and returned to the original plating media. All measurements on cultured neurons were taken between DIV 14 and DIV 18 (~9-14 d post transfection) to allow for sodium channel maturation (and thus spiking). No all-trans-retinal was supplemented for any cultured neuron recordings.

### Electrophysiology and fluorescence microscopy in cultured primary hippocampal neurons

Whole-cell patch clamp recordings of cultured neurons for **Supplementary Table 1** were acquired via an Axopatch 700B amplifier (Molecular Devices LLC) and Digidata 1440 digitizer (Molecular Devices LLC). Neurons were patched between DIV14 and DIV18 and were bathed in Tyrode’s solution (125mM NaCl, 2mM KCl, 3mM CaCl2, 1mM MgCl2, 10mM HEPES, 30mM glucose, pH 7.3 (NaOH adjusted)) at 32 °C during measurements. Synaptic blockers (NBQX, 10μM; d(-)-2-amino-5-phosphonovaleric acid, 25 μM; gabazine, 20μM; Tocris) were added to the extracellular solution for single-cell electrophysiology. Borosilicate glass pipettes with an outer diameter of 1mm and a wall thickness of 0.2mm were pulled to produce electrodes with resistance of 3-10 MΩ and were filled with an internal solution containing 135mM potassium gluconate, 8mM NaCl, 10mM HEPES, 4mM Mg-ATP, 0.4mM Na-GTP, 0.6mM MgCl2, 0.1mM CaCl2, pH 7.25 (KOH adjusted) at 295mOsm. Measurements from primary neuron cultures were performed on the electrophysiology setup described above. Patch-clamp data was acquired only if the resting potential was below −45mV and access resistance was <25 MΩ. Access resistance was compensated at 30-70%. Fluorescence imaging was performed on an inverted fluorescence microscope (Nikon Ti), equipped with a red laser (637nm, 100mV, Coherent, OBIS 637LX, Pigtailed) expanded by a beam expander (Thorlabs Inc) and focused onto the back focal plane of a 40×NA 1.15 objective lens (Nikon Corp.).

### *In utero* electroporation, AAV injection, and acute brain slice preparation

For IUE, embryonic day (E) 15.5 timed-pregnant female Swiss Webster (Taconic Biosciences Inc., Albany, NY) mice were deeply anesthetized with 2% isoflurane. Uterine horns were exposed and periodically rinsed with warm sterile PBS. Plasmid DNA (1-2 μg total at a final concentration of ~2-3 μg/μL diluted in sterile PBS) was injected into the lateral ventricle of one cerebral hemisphere of an embryo. Five voltage pulses (50 V, 50 ms duration, 1 Hz) were delivered using 5 mm round plate electrodes (ECM™ 830 electroporator, Harvard Apparatus), placing anode or cathode on the top of the skull to target cortex or hippocampus, respectively. Electroporated embryos were placed back into the dam, and allowed to mature to delivery. Brain slices were prepared from electroporated mice at P15-P22.

The electroporated mice were anaesthetized by isoflurane inhalation, decapitated, and cerebral hemispheres were quickly removed and placed in cold choline-based cutting solution consisting of (in mM): 110 choline chloride, 25 NaHCO3, 2.5 KCl, 7 MgCl2, 0.5 CaCl2, 1.25 NaH2PO4, 25 glucose, 11.6 ascorbic acid, and 3.1 pyruvic acid (339-341 mOsm/kg; pH 7.75 adjusted with NaOH) for 2 min, then blocked and transferred into a slicing chamber containing ice-cold choline-based cutting solution. Coronal slices (300 μm thick) were cut with a Compresstome VF-300 slicing machine, then transferred to a holding chamber with artificial cerebrospinal fluid (ACSF) containing (in mM) 125 NaCl, 2.5 KCl, 25 NaHCO3, 2 CaCl2, 1 MgCl2, 1.25 NaH2PO4 and 11 glucose (300-310 mOsm/kg; pH 7.35 adjusted with NaOH), and recovered for 10 min at 34 °C, followed by another 30 min at room temperature. Slices were subsequently maintained at room temperature until use. Both cutting solution and ACSF were constantly bubbled with 95% O2 and 5% CO2.

For AAV injection, 21-day-old C57 BL/6J mice were anesthetized with isoflurane and placed in a small animal stereotaxic apparatus (David Kopf Instruments, CA, USA). Animals were injected with a volume of 200 nl rAAV8-Syn-Archon1-KGC-EGFP-Kv2.1_motif_-ER2 using a Nanoject (Drummond Scientific Co Inc, Broomall, PA) via glass pipettes with 20-30 μm diameter tips in striatum: anteroposterior (AP) 1.2, mediolateral (ML) 2.1, dorsoventral (DV) 3.2. Brain slices were then prepared from these AAV-injected mice at postnatal day 30-35. Mice were deeply anesthetized with isoflurane and perfused transcardially using cold saline containing (in mM): 194 sucrose, 30 NaCl, 4.5 KCl, 1.2 NH_2_PO_4_, 0.2 CaCl_2_, 2 MgCl_2_, 26 NaHCOs, and 10 D-(+)-glucose saturated with 95% O_2_ and 5% CO_2_, pH=7.4 adjusted with NaOH, 320-340 mOsm/L. Coronal slices (250-300 μm thick) were cut using a slicer (VT1200 S, Leica Microsystems, USA) and then incubated for 10 – 15 min in a holding chamber (BSK4, Scientific System Design Inc., USA) at 32°C with regular ACSF containing (in mM): 136 NaCl, 3.5 KCl, 1 MgCl_2_, 2.5 CaCl_2_, 26 NaHCO_3_ and 11 glucose saturated with 95% O_2_ and 5% CO_2_, followed by at least one hour recovery at room temperature (21-25°C) before recording.

### Concurrent electrophysiology and fluorescence imaging in acute brain slice

For recording in **Figure 1** and **Supplementary Figure 7**, individual slices were transferred to a recording chamber mounted on an upright microscope (Olympus BX51WI, see below) and continuously superfused (2-3 mL/min) with carbogenated ACSF at room temperature. Whole cell patch-clamp recordings were performed with borosilicate glass pipettes (KG33, King Precision Glass Inc.) heat polished to obtain direct current resistances of ~4-6 MΩ. Pipettes were filled with an internal solution containing in mM: 120 K-Gluconate, 2 MgCl2, 10 HEPES, 0.5 EGTA, 0.2 Na2ATP, and 0.2 Na3GTP. Voltage clamp recordings were made with a microelectrode amplifier (Multiclamp 700B, Molecular Devices LLC). Cell membrane potential was held at −60 mV, unless specified otherwise. Signals were low-pass filtered at 2 kHz and sampled at 10-20 kHz with a Digidata 1440A (Molecular Devices LLC), and data were stored on a computer for subsequent off-line analysis. Cells in which the series resistance (Rs, typically 8-12 MΩ) changed by >20% were excluded from subsequent data analysis. In addition, cells with Rs more than 25 MΩ at any time during the recordings were discarded. In some cases, conventional characterization of neurons was made in both voltage and current clamp configurations. Positive neurons were identified for recordings on the basis of EGFP expression visualized with a microscope equipped with a standard GFP filter (BX-51WI, Olympus Corp.). Optical voltage recordings were taken through a 40x water immersion objective (Olympus LUMFL N 40x/0.8W). Fluorescence was excited using a fiber-coupled 637 nm red laser (140mW, Coherent Obis 637-140 LX), and the emission was filtered through a 664 long pass filter. Images were collected on an EMCCD camera (Andor iXON Ultra 888) or sCMOS camera (Andor Zyla4.2 Plus Andor) in a reduced pixel window to enable acquisition at ~1kHz. Each trial was about 30 seconds in duration.

For optical recordings in **Figure 3a, b, c** and **Supplementary Figure 1, 2, 3, 8**, acute brain slices were transferred to a recording chamber mounted on an inverted Eclipse Ti-E (Nikon) equipped with a CMOS camera (Zyla5.5, Andor), LEDs (Spectra, Lumencor), a 637nm Laser (637 LX, OBIS) focused on the back focal plane of a 40×NA 1.15 objective (Nikon), and a Polygon400 Multiwavelength Patterned Illuminator (Mightex) with 470 nm LED (ThorLabs), and continuously superfused (2-3 mL/min) with carbogenated ACSF at room temperature. Positive cells were imaged under 0.8 or 1.5 W/mm^2^ excitation light power at 637 nm from the laser. 4-aminopyridine at a final concentration of 1 mM was added to induce neuronal activity for experiments in **Supplementary Figure 2**. For **Figure 3a, b, c** and **Supplementary Figure 8** cells were illuminated with 2 ms blue light pulses at light power in the range from 0.1 to 1.0 mW/mm^2^.

### Mouse surgery

All *in vivo* mouse procedures were performed in accordance with the National Institute of Health Guide for Laboratory Animals and approved by the Boston University Institutional Animal Care and Use and Biosafety Committees.

#### Virus injection surgery

All AAV was produced by the University of North Carolina Chapel Hill Vector Core. Adult female C57BL/6 mice (Charles River Laboratories, Inc.), 8-12 weeks at the time of surgery, were used for all experiments. AAV-syn-SomArchon1 (5.9e12 genome copies (GC)/ml) or AAV-syn-SomArchon-P2A-CoChR-Kv2.1 (2.19e13 GC/ml) was injected into the motor cortex (AP: +1.5, ML: +/− 1.5, DV: −0.3, 0.5uL virus), visual cortex (AP: −3.6, ML: +/−2.5, DV: −0.3, 0.5uL virus), hippocampus (AP:-2.0, ML:+1.4, DV:-1.6, 1uL virus) or striatum (AP:+0.8, ML:-1.8, DV:-2.1, 1uL virus). Viral injection occurred at 50-100nL/min (ten minutes total) using a 10uL syringe (NANOFIL, World Precision Instruments LLC) fitted with a 33 gauge needle (World Precision Instruments LLC, NF33BL) and controlled by a microinfusion pump (World Precision Instruments LLC, UltraMicroPump3-4). The syringe was left in place for an additional 10 minutes following injection to facilitate viral spread. About one week following the viral injection, mice underwent a second surgery to implant the cranial window for *in vivo* imaging.

#### Cortex imaging window implantation

The imaging window consisted of a stainless steel cannula (OD: 3.17mm, ID: 2.36mm, 1mm height, AmazonSupply, B004TUE45E) fitted with a circular coverslip (#0, OD: 3mm, Deckgläser Cover Glasses, Warner Instruments LLC, 64-0726 (CS-3R-0)) adhered using a UV curable glue (Norland Products Inc., Norland Optical Adhesive 60, P/N 6001). A craniotomy of ~3mm in diameter was created, with the dura left intact, over the motor cortex (centered at AP: +1.5, ML: +/−1.75) or visual cortex (AP: −3.6, ML: +/− 2.15). The imaging window was positioned over the cortex so that it was flush with the dura surface. Kwik sil adhesive (World Precision Instruments LLC, KWIK-SIL) was applied around the edges of the imaging window to hold the imaging window in place and to prevent any dental cement from touching the brain. Three small screws (J.I. Morris Co., F000CE094) were screwed into the skull to further anchor the imaging window to the skull. Dental cement was then gently applied to affix the imaging window to the exposed skull, and to mount an aluminum headbar posterior to the imaging window.

#### Hippocampus and striatum imaging window implantation

Hippocampal and striatal window surgeries were performed similar to those described previously^48^. For each imaging window, a virus/drug infusion cannula (26G, PlasticsOne Inc., C135GS-4/SPC) was attached to a stainless steel imaging cannula (OD: 3.17mm, ID: 2.36mm, 1 or 2mm height, AmazonSupply, B004TUE45E). The bottom of the infusion cannula was flush with the base of the stainless steel cannula, and a circular coverslip (#0, OD: 3mm, Deckgläser Cover Glasses, Warner Instruments Inc., 64-0726 (CS-3R-0)) was adhered using a UV curable glue (Norland Products Inc., Norland Optical Adhesive 60, P/N 6001). An additional insulated stainless steel wire (Diameter: 130um, PlasticsOne Inc., 005SW-30S, 7N003736501F) was glued to the viral/drug infusion cannula with super glue (Henkel Corp., Loctite 414 and Loctite 713) and protruded from the bottom of the infusion cannula and imaging window by about 200um for LFP recordings.

A craniotomy ~3mm in diameter was made over the hippocampus CA1 region (AP: −2.0, ML: +2.0) or the striatum (AP: +0.8, ML: −1.8). A small notch was made on the posterior edge of the craniotomy to accommodate the infusion cannula and LFP recording electrode. The overlying cortex was gently aspirated using the corpus callosum as a landmark. The corpus callosum was then carefully thinned in order to expose the hippocampus CA1 region or the dorsal striatum. The imaging window was positioned in the craniotomy, and Kwik sil adhesive (World Precision Instruments LLC, KWIK-SIL) was applied around the edges of the imaging window to hold it in place and to prevent any dental cement from touching the brain. Three small screws (J.I. Morris Co., F000CE094) were screwed into the skull to further anchor the imaging window to the skull, and a small ground pin was inserted into the posterior part of the brain near the lambda suture as a ground reference for LFP recordings. Dental cement was then gently applied to affix the imaging window to the exposed skull, and to mount an aluminum headbar posterior to the imaging window.

In mice that did not receive a virus injection prior to window implantation, 1uL of AAV-syn-SomArchon (5.9e12 GC/ml) or 1uL of AAV-syn-SomArchon-P2A-CoChR-Kv2.1 (2.19e13 GC/ml) was injected through the virus/drug infusion cannula at 100nL/min through an internal infusion cannula (33G, PlasticsOne Inc., C315IS-4/SPC) connected to a microinfusion pump (World Precision Instruments LLC, UltraMicroPump3-4), one week after the window implantation surgery. The internal infusion cannula was left in place for 10 minutes following injection to facilitate viral spread. Mice were awake and head-fixed throughout the injection period.

All mice were treated with buprenex for 48 hours following surgery and single-housed to prevent any damage to the headbar or window implant.

### *In vivo* imaging in the live mouse brain

All optical recordings were acquired on a conventional one-photon fluorescence microscope equipped with an Orca Flash 4.0 V3 Digital CMOS camera (Hamamatsu Photonics K.K., C13440-20CU), 10x NA0.25 LMPlanFI air objective (Olympus Corp.), 40x NA0.8 LUMPlanFI/IR water immersion objective (Olympus Corp.), 20x NA1.0 XLUMPlanFL N water immersion objective (Olympus Corp.), 470nm LED (ThorLabs Inc., M470L3), 140mW 637nm red laser (Coherent Obis 637-140X), a green filter set with a 470/25nm bandpass excitation filter, a 495nm dichroic, and a 525/50nm bandpass emission filter, and a near infrared filter set with a 635nm laser dichroic filter, and a 664nm long pass emission filter. The near infrared laser illuminated an area of ~80-120um in diameter, and a mechanical shutter (Newport Corporation, model 76995) was positioned in the laser path to control the timing of illumination over the imaging window. Optical recordings were acquired at 390-900Hz with HCImage Live (Hamamatsu Photonics K.K.) or NIS Elements (Nikon Instruments) software. HC Image Live data were stored as DCAM image files (DCIMG), and further analyzed offline in Fiji/ImageJ and Matlab (Mathworks Inc.). NIS Elements data were stored as.nd2 files and further analyzed offline using the NIS Elements software.

The GFP signal of SomArchon was acquired in the green channel (λ_ex_=470nm) at 1024×1024 pixels with 2×2 binning to show cell structure and distribution. Optical voltage recordings were imaged in the near infrared channel (λ_ex_=637nm) with 2×2 or 4×4 binning. OmniPlex system (PLEXON Inc.) was used to synchronize data acquisition from different systems. In all experiments, the OmniPlex system recorded the start of image acquisition from the sCMOS camera, the acquisition time of each frame, and other experiment-dependent signals described below.

#### Optical imaging of spontaneous neural activity

All *in vivo* optical imaging of spontaneous neural activity was performed when mice were awake and head fixed in a custom holder that allowed for attachment of the headplate at the anterior end. Animals were covered with an elastic wrap to prevent upward movement. Spontaneous neural activity recordings lasted continuously up to 30,000 frames (~36 seconds).

#### Eye puff

During some *in vivo* hippocampal imaging recordings, an eye puff was applied to evoke high frequency local field potential responses in the hippocampus (**Supplementary Figure 14**). The mice were head fixed in a custom holder that allowed for attachment of the headplate at the anterior end, and they were covered with an elastic wrap to prevent upward movement. Each experimental session consisted of 20-30 trials, with each trial lasting for 5000 frames (~6 seconds). Three seconds after the start of image acquisition, the sCMOS camera sent a TTL pulse to a function generator (Agilent Technologies, model 33210A), which triggered a 100ms long air puff. The air puff was 5-10 psi, and administered via a 0.5mm cannula placed 2-3cm away from the mouse’s eye. The puff TTL pulses were also recorded with the OmniPlex system (PLEXON Inc.). Eye movement was monitored using a USB webcam (Logitech, Carl Zeiss Tessar 2.0/3.7 2MP Autofocus).

#### Optopatch blue light stimulation

All *in vivo* optopatch experiments were performed when mice were awake and head fixed in a custom holder that allowed for attachment of the headplate at the anterior end. Animals were covered with an elastic wrap to prevent upward movement. A 470nm LED (ThorLabs Inc., M470L3) was coupled to a Polygon400 Multiwavelength Patterned Illuminator (Mightex), and the blue light was focused through the objective lens to illuminate the center of the field of view. At the onset of imaging, the sCMOS camera sent a TTL pulse to trigger Axon CNS (Molecular Devices LLC, Digidata 1440A) which controlled the 470 nm LED (ThorLabs). Each trial lasted 1.1 second and consisted of a single 100 ms long blue light pulse, 500 ms after trial onset. Each recording session consisted of 10 trials with increasing blue light power from 0.1 to 1 mW/mm^2^, with a step of ~ 0.1 mW/mm^2^ per trial. The OmniPlex system (PLEXON Inc.) recorded the timing of TTL pulses used to trigger the Axon CNS.

#### Head-fixed voluntary movement experiments

All voluntary movement experiments were performed while awake, head-fixed mice were freely navigating a spherical treadmill. The spherical treadmill was constructed following the design of Dombeck et al. 2007^49^. Briefly, a 3D spherical Styrofoam ball was supported by air, and motion was tracked using two computer mouse sensors positioned roughly +/− 45 degrees from center along the equator of the ball. All motion sensor displacement data was acquired at 100 Hz on a separate computer and synthesized using a custom Python script. Motion sensor displacement data were then sent to the image acquisition computer to be accumulated using a modified ViRMEn Matlab script. The timing of each motion sensor displacement data point was also recorded using the OmniPlex system (PLEXON Inc.) to synchronize movement data with optical voltage recordings.

In order to determine the mouse movement speed, ball movement was first calibrated. The ball was pinned on the two sides, and rotated vertically to calibrate sensor displacement.

All mice were habituated on the spherical treadmill for at least three days, at least 20 minutes per day, prior to image acquisition. During optical imaging, mice were imaged while freely navigating the spherical treadmill. Each field of view was recorded for at least 36 seconds total. In some fields of view, we performed multiple trials, and each trial was at least 12 seconds in duration with an inter-trial interval of at least 30 seconds in duration.

### Local field potential recording

Local field potentials were recorded using an OmniPlex system (PLEXON Inc.) at a 1 kHz sampling rate. To synchronize optical recordings with LFP recordings, the camera sent out a TTL pulse to the OmniPlex system at the onset of imaging and after each acquired frame.

### Motion correction

In **Figures 2i, 4**, and **5**, motion correction was performed with a custom Python script. For each field of view (FOV), if multiple video imaging files were collected for the same FOV, we started with the first imaging file to ensure speedy data processing (a single video file contains a series of images). We first generated the reference image by averaging across all images within the file. We then performed a series of image processing procedures to enhance the contrast of the reference image and every image in the file to facilitate motion correction. We first removed 10% of the pixels along all edges of an image to remove any camera induced artifact. We then applied a high-pass filter (Python scipy package, ndimage.gaussian_filter, sigma=50) to remove low frequency components within the images. To enhance the boundary of high intensity areas, we identified the boundary as the difference between two low pass filtered images (sigma=2 and 1). We then enhanced the boundary by adding 100 times the boundary back to the low pass filtered image (sigma=2). We then limited the intensity range of the processed images within one standard deviation above and below the average intensity of the image, by setting the pixels with intensity higher than mean+std as mean+std, and the pixels with intensity lower than mean-std as zeroes. Finally, to counter any potential bleaching over time, we normalized the intensity of each image by shifting the mean intensity to zero and divided intensity values by the standard deviation of all pixel intensities in that image. After image processing, we calculated the displacement of each image, by identifying the max cross-correlation coefficient between each image and the reference image, and then corrected motion by shifting the displacement in the original, unenhanced image sequence. If the same FOV was imaged over an extended period of time, where multiple files were acquired, we motion-corrected subsequent files by aligning them to the first file, so that the same ROIs from the same FOV could be applied across the entire imaging session. Specifically, we first refined the reference frame by generating the mean intensity projection image from the motion-corrected first imaging file. The refined reference image was then used to motion-correct all files of the same FOV, including the first file, using the same procedure described above. The motion-corrected, original, unenhanced image sequences, were then used for subsequent manual ROI segmentation, and further analysis.

### ROI identification

We imported the image files (motion-corrected as above, if necessary) into Fiji/ImageJ or NIS Elements and manually segmented ROIs by examining the time-series images to identify the area with clear neuron outlines and/or intensity dynamics over time. The optically-recorded voltage traces for each ROI were generated from the motion-corrected image sequences using the “multiple measurement” function and were then used for analyses.

For **Figure 2l** and **Figure 6**, cells were densely packed, so we identified and tracked ROIs semi-manually across image sequences without performing motion correction. We first visually inspected all image sequences and identified those with minimal motion and with an SNR greater than ~2 for further analysis. We then performed an iterative ROI-selection procedure to identify ROIs that best fit each cell. Specifically, we started by manually selecting ROIs from the max projection image of the entire image sequence. The image sequence was then visually inspected to identify frames whose cells exhibited shifts of more than three pixels from the defined ROI. We then used these frames to separate the image sequence into multiple time intervals, and obtained a new set of max projection images to identify new ROIs within these time intervals for these cells. This procedure was repeated iteratively until the ROI represented the cell across all image frames in their corresponding time intervals without the cell moving out of the ROI. Thus, with this procedure, we created multiple ROIs representing the same cell across different frames. For each cell, we extracted traces for every ROI during its corresponding time interval, and stitched the baseline-normalized traces for the same cell(s) in time. The fluorescence traces of each cell were then detrended for further analysis. See **Supplementary Figure 15** for an example of raw and processed traces for two cells in the same field of view.

### Hippocampal spike detection

Spikes were associated with a rapid increase in intensity, followed by a rapid decrease. In contrast, occasional motion artifacts were usually associated with a decrease in intensity as a neuron moved out of the ROI. To facilitate spike detection, we first removed motion artifacts. For each time point of the fluorescence intensity trace for each ROI, we calculated the change in intensity from that of the prior time point (I_change_). We then defined noise as the time points where instantaneous I_change_ were three standard deviations below the mean value of I_change_ across the entire trace. We excluded any time points where the I_change_ of the previous time point was more than one standard deviation above the average I_change_, because this might have indicated a spike. These noise time points and their following three time points (since we found that motion artifacts are typically >4ms) were then considered motion artifacts, and removed from further analysis. We then recalculated the standard deviation of the I_change_, excluding the data points of motion artifact. The peaks of spikes were then identified as time points with the following two criteria: 1) the intensity change of the time point combined with that of its preceding time point was more than three standard deviations above the average I_change_, and 2) the intensity change over the next two time points was less than two standard deviations below the average I_change_.

### Hippocampal spike phase calculation

Hippocampal spike-phase analysis was performed on 16 neurons from 7 FOVs from 4 mice. For each FOV, we analyzed data collected over 10 trials (~60 seconds total), where animals experienced an eye puff in each trial as described above in *Eye puff*. To calculate the phase of spikes at theta frequency (4-10 Hz), we first band-pass filtered both the optical voltage trace and the simultaneously recorded LFP at theta frequency (eegfilt, EEGLAB toolbox). The peaks of theta oscillation power were then identified with the findspike function in Matlab. For each spike, we obtained the phase of the spike by calculating the timing of each spike relative to the period of that oscillation cycle in degrees. We averaged the phases of all spikes from the same neuron as the average phase of a given neuron.

### Analysis for pair-wise coherence between hippocampal neurons and LFPs

The coherence analysis was performed on 9 FOVs that contained multiple neurons from 4 mice. Each FOV contained imaging data over a period of 6-36 seconds. For each FOV, we first re-sampled the LFP at the acquisition rate of the optical imaging. We then divided the optical voltage traces and LFPs into segments of 1000 data points. We then calculated the averaged coherence, at theta frequency (4-10 Hz), with the functions in the Chronux toolbox (optical voltage trace to optical voltage trace or LFP: coherencyc, and spike to spike: coherencypt) with tapers=[10 19], fpass=[4 10] and trialave=1. To compare V_mo_-V_mo_ coherence vs. V_mo_-LFP coherence across nine FOVs, we averaged the coherence of neurons in the same FOV to obtain the mean coherence of that FOV, and then performed statistical tests across FOVs using the individual FOV’s mean coherences. To understand the relationships between pairs of coherence, we used the Matlab function, fitlm, to perform a linear regression between coherent pairs and obtain the p-value and r^2^ value.

### Striatum, motor cortex, and visual cortex, spike detection

After motion correction, we first identified large fluorescence increases using a threshold of 4 standard deviations above the baseline. The baseline was manually selected as a period of >500ms without spiking or drifting due to z-plane shifting or photobleaching. From these large fluorescence increases, we selected those with shorter than 4 ms rise times and 4 ms decay times as spikes.

### Firing rate comparison of striatal neurons during high versus low speed movement

Animals’ movement data was first interpolated to the voltage imaging frame rate with Matlab function interp1, and then smoothed using a 1.5Hz low pass Butterworth filter to remove any motion sensor artifact. We calculated the average movement speed at 0.5-second intervals and defined low speed periods as intervals where the average speed was ≤5cm/s and high speed periods as intervals where the average speed was ≥10cm/s. The firing rates during these high and low motion periods were compared, and a two-sided Wilcoxon rank sum test was used to determine significance between these periods.

### Detrending

All optically-recorded SomArchon traces reported in the manuscript (except those in **Figure 5**) were corrected for photobleaching or focus shift by subtracting baseline fluorescence traces that were low-pass filtered and fit to a double or single exponential function.

### Histology

Mice were transcardially perfused with PBS followed by 4% paraformaldehyde. The brain was gently extracted from the skull and post-fixed in 4% paraformaldehyde at room temperature for 1-4 hours. Fixed brains were transferred to a 30% sucrose-PBS solution and rotated 24-48 hours at 4C for cryoprotection. Cryoprotected brains were frozen in OCT in a dry ice bath and sliced (coronal) to 50um thickness using a cryostat.

### Sample size

No statistical methods were used to estimate sample size for animal studies throughout. We did not perform a power analysis, since our goal was to create a new technology; in the reference (Dell, R. B., Holleran, S. & Ramakrishnan, R. Sample size determination. ILAR. J. 43, 207-213 (2002)), as recommended by the NIH, “In experiments based on the success or failure of a desired goal, the number of animals required is difficult to estimate…” As noted in the aforementioned paper, “The number of animals required is usually estimated by experience instead of by any formal statistical calculation, although the procedures will be terminated [when the goal is achieved].” These numbers reflect our past experience in developing neurotechnologies.

### Data exclusions

Voltage imaging datasets with significant motion or where no spikes were detected were excluded from analysis.

### Replication

All attempts at replication were successful.

### Randomization and blinding

No randomization or blinding were used for animal studies throughout.

## Acknowledgments

We thank Tina Ta for her help on histology. We thank M. Murdock for help with animal work, and N. Pak for help with assembling the *in vivo* imaging setup.

## Author Contributions

K.D.P. and E.S.B. initiated the project. K.D.P., S.B., H.T., S.N.S., X.H., and E.S.B. designed all *in vivo* experiments and interpreted the data. K.D.P. developed SomArchon and together with E.E.J., O.A.S., and E.C. characterized all constructs in cultured cells. K.D.P., V.G.L.H., D.P., C.S., and B.L.S. performed characterization of SomArchon in acute brain slices. S.B., S.N.S. and H.J.G. performed all mouse surgeries for *in vivo* experiments. M.F.R. assisted on imaging setups. K.D.P, S.B., H.T., and S.N.S performed all *in vivo* imaging experiments and analyzed all *in vivo* imaging data. K.D.P., S.B., H.T., S.N.S, X.H. and E.S.B. wrote the paper with contributions from all of the authors. E.S.B. and X.H. oversaw all aspects of the project. The funders had no role in study design, data collection and analysis, decision to publish, or preparation of the manuscript.

## Competing interests

The authors declare no competing financial interests.

## Funding

E.S.B. acknowledges funding from Edward and Kay Poitras, NIH Director’s Pioneer Award 1DP1NS087724, NIH 1R01GM104948, NIH 1R01EB024261, NIH 1R01DA045549, NIH 1R01MH114031, Charles Hieken, NIH 1R01NS102727, John Doerr, NSF Grant 1734870, the HHMI-Simons Faculty Scholars Program, Human Frontier Science Program RGP0015/2016, NIH 1R43MH109332, U. S. Army Research Laboratory and the U. S. Army Research Office under contract/grant number W911NF1510548, NIH 2R01DA029639, and NSF CBET 1344219. X.H. acknowledges funding from NIH Director’s Office (1DP2NS082126), NINDS (1R01NS081716, 1R01NS087950-01), the Grainger Foundation, the Pew Foundation, Boston University Biomedical Engineering Department. S.B. and S.N.S. acknowledge funding from the NIH/NIGMS T32 Quantitative Biology and Physiology Fellowship (GM008764) through the Boston University Biomedical Engineering department. The funders had no role in study design, data collection and analysis, decision to publish, or preparation of the manuscript.

## Code availability

Computer code used to generate results for this study are available from the corresponding author upon reasonable request.

## Data availability

The data that support the findings of this study are available from the corresponding author upon reasonable request. Sequences of the reported proteins will be deposited at Genbank upon acceptance.

