## Supplementary Materials for "Population imaging of neural activity in awake behaving mice in multiple brain regions"

### Supplementary Information

**Supplementary Table 1.** Characteristics of screened candidates of soma-localized Archon1 voltage sensor.

| **Full name of construct** | **How far from the soma, approximately, was the fluorescence detected using visual inspection, in this study?** | **Membrane/cytoplasmic localization detected using visual inspection in near-infrared channel, in this study** | **Voltage sensitivity*** |
|---|---|---|---|
| Archon1-linker-KA2(1-150)-EGFP | 50-100 μm | Cytoplasmic with bright puncta in soma | Non-functional |
| Archon1-linker-KA2(1-100)-EGFP | 50-100 μm | Cytoplasmic with bright puncta in soma | Non-functional |
| KA2(1-150)-linker-Archon1-EGFP | 50-100 μm | Cytoplasmic with bright puncta in soma | Non-functional |
| Archon1-KGC-EGFP-NaV1.2(II-III)-ER2 | 80-100 μm | Membrane with significant aggregation in soma | Not measured |
| Archon1-KGC-EGFP-NaV1.6(II-III)-ER2 | 30-60 μm | Membrane with no aggregation | 12% of ΔF/F per 100 mV voltage step |
| Archon1-KGC-EGFP-Kv2.1-motif-ER2 | 20-40 μm | Membrane with no aggregation | 30% of ΔF/F per 100 mV voltage step |
| Archon1-KGC-EGFP-AnkMB(270)-motif-ER2 | >100 μm | Cytoplasmic with bright puncta in soma | Non-functional |
| Archon1-KGC-EGFP-AnkMB(490)-motif-ER2 | >100 μm | Cytoplasmic with bright puncta in soma | Non-functional |
| Archon1-KGC-EGFP-AnkSB-motif-ER2 | >100 μm | Cytoplasmic with bright puncta in soma | Non-functional |
| Archon1-KGC-EGFP-AnkCt-motif-ER2 | 50-70 μm | Membrane with minor aggregation in soma | 15% of ΔF/F per 100 mV voltage step |
| Archon1-KGC-EGFP-AnkSR-motif-ER2 | 40-60 μm | Membrane with minor aggregation in soma | 15% of ΔF/F per 100 mV voltage step |
| Archon1-KGC-EGFP-AnkTail-motif-ER2 | >100 μm | Cytoplasmic with bright puncta in soma | Non-functional |

*voltage sensitivity was quantified by whole-cell patch clamp.

Amino acid sequences:

**Linker:**
GGSGGTGGSGGT

**KA2(1-150)**
MPAELLLLLIVAFANPSCQVLSSLRMAAILDDQTVCGRGERLALALAREQINGIIEVPAK
ARVEVDIFELQRDSQYETTDTMCQILPKGVVSVLGPSSSPASASTVSHICGEKEIPHIKVG
PEETPRLQYLRFASVSLYPSNEDVSLAVS

**KA2(1-100)**
MPAELLLLLIVAFANPSCQVLSSLRMAAILDDQTVCGRGERLALALAREQINGIIEVPAK
ARVEVDIFELQRDSQYETTDTMCQILPKGVVSVLGPSSSP

**KGC:**
KSRITSEGEYIPLDQIDINV

**ER2:**
FCYENEV

**NaV1.2(II-III):**
SSFSSDNLAATDDDNEMNNLQIAVGRMQKGIDFVKRKIREFIQKAFVRKQKALDEIKPL
EDLNNKKDSCISNHTTIEIGKDLNYLKDGNGTTSGIGSSVEKYVVDESDYMSFINNPSLT
VTVPIALGESDFENLNTEEFSSESDMEESKEKLNATSSSEGSTVDIGAPAEGEQPEAEPEE
SLEPEACFTEDCVRKFKCCQISIEEGKGKLWWNLRKTCYKIS

**NaV1.6(II-III)**
TVRVPIAVGESDFENLNTEDVSSESDP

**Kv2.1-motif:**
QSQPILNTKEMAPQSKPPEELEMSSMPSPVAPLPARTEGVIDMRSMSSIDSFISCATDFPE
ATRF

**AnkMB(270)-motif (270-kDa AnkyrinG (1-837aa)):**
MAHAASQLKKNRDLEINAEEETEKKKKHRKRSRDRKKKSDANASYLRAARAGHLEKA
LDYIKNGVDVNICNQNGLNALHLASKEGHVEVVSELLQREANVDAATKKGNTALHIAS
LAGQAEVVKVLVTNGANVNAQSQNGFTPLYMAAQENHLEVVRFLLDNGASQSLATED
GFTPLAVALQQGHDQVVSLLLENDTKGKVRLPALHIAARKDDTKAAALLLQNDTNADI
ESKMVVNRATESGFTSLHIAAHYGNINVATLLLNRAAAVDFTARNDITPLHVASKRGNA
NMVKLLLDRGAKIDAKTRDGLTPLHCGARSGHEQVVEMLLDRAAPILSKTKNGLSPLH
MATQGDHLNCVQLLLQHNVPVDDVTNDYLTALHVAAHCGHYKVAKVLLDKKANPNA
KALNGFTPLHIACKKNRIRVMELLLKHGASIQAVTESGLTPIHVAAFMGHVNIVSQLMH
HGASPNTTNVRGETALHMAARSGQAEVVRYLVQDGAQVEAKAKDDQTPLHISARLGK
ADIVQQLLQQGASPNAATTSGYTPLHLSAREGHEDVAAFLLDHGASLSITTKKGFTPLH
VAAKYGKLEVASLLLQKSASPDAAGKSGLTPLHVAAHYDNQKVALLLLDQGASPHAA
AKNGYTPLHIAAKKNQMDIATSLLEYGADANAVTRQGIASVHLAAQEGHVDMVSLLLS

RNANVNLSNKSGLTPLHLAAQEDRVNVAEVLVNQGAHVDAQTKMGYTPLHVGCHYG
NIKIVNFLLQHSAKVNAKTKNGYTPLHQAAQQGHTHIINVLLQNNASPNELTVNGNTAL
AIARRLGYISVVDTLKVVTEEIMTTTTIT

**AnkMB(490)-motif (490-kDa AnkyrinG (1-800aa))**
MAHAASQLKKNRDLEINAEEETEKKRKHRKRSRDRKKKSDANASYLRAARAGHLEKA
LDYIKNGVDVNICNQNGLNALHLASKEGHVEVVSELLQREANVDAATKKGNTALHIAS
LAGQAEVVKVLVTNGANVNAQSQNGFTPLYMAAQENHLEVVRFLLDNGASQSLATED
GFTPLAVALQQGHDQVVSLLLENDTKGKVRLPALHIAARKDDTKAAALLLQNDTNAD
VESKSGFTPLHIAAHYGNINVATLLLNRAAAVDFTARNDITPLHVASKRGNANMVKLLL
DRGAKIDAKTRDGLTPLHCGARSGHEQVVEMLLDRSAPILSKTKNGLSPLHMATQGDH
LNCVQLLLQHNVPVDDVTNDYLTALHVAAHCGHYKVAKVLLDKKASPNAKALNGFTP
LHIACKKNRIRVMELLLKHGASIQAVTESGLTPIHVAAFMGHVNIVSQLMHHGASPNTT
NVRGETALHMAARSGQAEVVRYLVQDGAQVEAKAKDDQTPLHISARLGKADIVQQLL
QQGASPNAATTSGYTPLHLAAREGHEDVAAFLLDHGASLSITTKKGFTPLHVAAKYGKL
EVASLLLQKSASPDAAGKSGLTPLHVAAHYDNQKVALLLLDQGASPHAAAKNGYTPLH
IAAKKNQMDIATSLLEYGADANAVTRQGIASVHLAAQEGHVDMVSLLLSRNANVNLSN
KSGLTPLHLAAQEDRVNVAEVLVNQGAHVDAQTKMGYTPLHVGCHYGNIKIVNFLLQ
HSAKVNAKTKNGYTALHQAAQQGHTHIINVLLQNNASPNELTVNGNTAL

**AnkSB-motif (270-kDa AnkyrinG (801-1521aa)):**
AIARRLGYISVVDTLKVVTEEIMTTTTITEKHKMNVPETMNEVLDMSDDEVRKASAPEK
LSDGEYISDGEEGEDAITGDTDKYLGPQDLKELGDDSLPAEGYVGFSLGARSASLRSFSS
DRSYTLNRSSYARDSMMIEELLVPSKEQHLTFTREFDSDSLRHYSWAADTLDNVNLVSS
PVHSGFLVSFMVDARGGSMRGSRHHGMRIIIPPRKCTAPTRITCRLVKRHKLANPPPMVE
GEGLASRLVEMGPAGAQFLGPVIVEIPHFGSMRGKERELIVLRSENGETWKEHQFDSKN
EDLAELLNGMDEELDSPEELGTKRICRIITKDFPQYFAVVSRIKQESNQIGPEGGILSSTTV
PLVQASFPEGALTKRIRVGLQAQPVPEETVKKILGNKATFSPIVTVEPRRRKFHKPITMTI
PVPPPSGEGVSNGYKGDATPNLRLLCSITGGTSPAQWEDITGTTPLTFIKDCVSFTTNVSA
RFWLADCHQVLETVGLASQLYRELICVPYMAKFVVFAKTNDPVESSLRCFCMTDDRVD
KTLEQQENFEEVARSKDIEVLEGKPIYVDCYGNLAPLTKGGQQLVFNFYSFKENRLPFSI
KIRDTSQEPCGRLSFLKEPKTTKGLPQTAVCNLNITLPAHKKETESDQDDAEKADRRQSF
ASLALRKRYSYLTEPSMKTVERSSGTARSLPTTYSHKPFFSTRPYQSWTTAPITVPGPAKS
GSLSSSPSNTPSA

**AnkCt-motif (270-kDa AnkyrinG (2334-2622aa))**
RTDIRMAIVADHLGLSWTELARELNFSVDEINQIRVENPNSLISQSFMLLKKWVTRDGKN
ATTDALTSVLTKINRIDIVTLLEGPIFDYGNISGTRSFADENNVFHDPVDGWQNETPSGSL
ESPAQARRLTGGLLDRLDDSSDQARDSITSYLTGEPGKIEANGNHTAEVIPEAKAKPYFP
ESQNDIGKQSIKENLKPKTHGCGRTEEPVSPLTAYQKSLEETSKLVIEDAPKPCVPVGMK
KMTRTTADGKARLNLQEEEGSTRSEPKQGEGYKVKTKKEIRNVEKKTH

**AnkSR-motif (270-kDa AnkyrinG (1534 -1933aa))**
SPLKSIWSVSTPSPIKSTLGASTTSSVKSISDVASPIRSFRTVSSPIKTVVSPSPYNPQVASGT
LGRVPTITEATPIKGLAPNSTFSSRTSPVTTAGSLLERSSITMTPPASPKSNITMYSSSLPFK
SIITSATPLISSPLKSVVSPTKSAADVISTAKATMASSLSSPLKQMSGHAEVALVNGSVSPL
KYPSSSALINGCKATATLQDKISTATNAVSSVVSAASDTVEKALSTTTAMPFSPLRSYVS
AAPSAFQSLRTPSASALYTSLGSSIAATTSSVTSSIITVPVYSVVNVLPEPALKKLPDSNSF
TKSAAALLSPIKTLTTETRPQPHFNRTSSPVKSSLFLASSALKPSVPSSLSSSQEILKDVAE
MKEDLMRMTAILQTDVPEEKPFQTDLP

**AnkTail-motif (270-kDa AnkyrinG (1934-2333aa)):**
REGRIDDEEPFKIVEKVKEDLVKVSEILKKDVCVESKGPPKSPKSDKGHSPEDDWTEFSS
EEIREARQAAASHAPSLPERVHGKANLTRVIDYLTNDIGSSSLTNLKYKFEEAKKDGEER
QKRILKPAMALQEHKLKMPPASMRPSTSEKELCKMADSFFGADAILESPDDFSQHDQDK
SPLSDSGFETRSEKTPSAPQSAESTGPKPLFHEVPIPPVITETRTEVVHVIRSYEPSSGEIPQS
QPEDPVSPKPSPTFMELEPKPTTSSIKEKVKAFQMKASSEEEDHSRVLSKGMRVKEETHI
TTTTRMVYHSPPGGECASERIEETMSVHDIMKAFQSGRDPSKELAGLFEHKSAMSPDVA
KSAAETSAQHAEKDSQMKPKLERIIEVHIEKGPQSPCE

**Supplementary Table 2**. Statistical analyses for **Figure 1d**, **Figure 4f**, **Figure 5, Figure 6,** and **Supplementary Figures 4, 7, 8**

Two-sided Wilcoxon Rank Sum test for **Figure 1d** between electrical and optical FWHM (full width, half maximum) of SomArchon in intact mouse brain slices

| **FWHM** | **Cortex** |
|---|---|
| p-value | 0.0010 |
| Wilcoxon rank sum statistic | 131 |

| **FWHM** | **Hippocampus** |
|---|---|
| p-value | 0.0019 |
| Wilcoxon rank sum statistic | 40 |

| **FWHM** | **Striatum** |
|---|---|
| p-value | 0.0411 |
| Wilcoxon rank sum statistic | 26 |

Two-sided Wilcoxon Rank Sum test for **Figure 4f** comparing firing rate during periods of low speed movement vs. high speed movement

| **Cell** | **p-value** | **Wilcoxon rank sum statistic** | **zval** |
|---|---|---|---|
| 1 | 0.583 | 698.5 | 0.5497 |
| 2 | 0.072 | 451 | -1.7989 |
| 3 | 0.043 | 417.5 | -2.0238 |
| 4 | 2.27E-07 | 3303 | -5.176 |
| 5 | 7.23E-05 | 2359 | 3.9684 |
| 6 | 0.525 | 1071 | -0.6359 |
| 7 | 0.655 | 1086 | -0.4468 |
| 8 | 0.0012 | 865.5 | -3.228 |
| 9 | 0.815 | 1084.5 | 0.2336 |
| 10 | 0.219 | 1418.5 | 1.228 |
| 11 | 0.144 | 1075 | -1.4617 |
| 12 | 0.0015 | 7239 | 3.1753 |
| 13 | 0.782 | 1296.5 | -0.2768 |
| 14 | 1.32E-04 | -3.8227 | 920.5 |

$\chi^2$ test for neurons used in **Figure 5** testing against the null hypothesis of a uniform distribution. DOF is degrees of freedom.

| | **LFP** | | | **$V_{mo}$** | | |
|---|---|---|---|---|---|---|
| **Cell #** | **p-value** | **$\chi^2$ test statistic** | **DOF** | **p-value** | **$\chi^2$ test statistic** | **DOF** |
| 1 | 1.58E-07 | 64.925 | 17 | 1.10E-07 | 65.849 | 17 |
| 2 | 0.0557 | 27.168 | 17 | 6.23E-40 | 232.946 | 17 |
| 3 | 0.619 | 14.670 | 17 | 2.23E-29 | 180.564 | 17 |
| 4 | 0.00117 | 40.308 | 17 | 4.14E-171 | 856.475 | 17 |
| 5 | 0.12 | 23.983 | 17 | 1.67E-49 | 279.756 | 17 |
| 6 | 0.51 | 16.190 | 17 | 1.72E-31 | 191.132 | 17 |
| 7 | 0.271 | 20.064 | 17 | 3.66E-20 | 133.677 | 17 |
| 8 | 0.618 | 14.688 | 17 | 1.11E-04 | 47.280 | 17 |
| 9 | 0.643 | 14.340 | 17 | 7.73E-13 | 94.909 | 17 |
| 10 | 0.718 | 13.263 | 17 | 3.92E-02 | 28.526 | 17 |
| 11 | 0.0263 | 30.000 | 17 | 1.14E-08 | 71.596 | 17 |
| 12 | 4.54E-06 | 56.109 | 17 | 1.00E-10 | 83.263 | 17 |
| 13 | 2.23E-34 | 205.508 | 17 | 2.40E-165 | 829.457 | 17 |
| 14 | 0.0636 | 26.641 | 17 | 5.47E-02 | 27.237 | 17 |
| 15 | 2.87E-05 | 51.073 | 17 | 1.46E-22 | 146.035 | 17 |
| 16 | 0.891 | 10.283 | 17 | 4.57E-04 | 43.146 | 17 |

Two-sided Wilcoxon Rank Sum test between the averaged $V_{mo}$-$V_{mo}$ coherence and $V_{mo}$-LFP coherence in main text results about **Figure 6, "Population imaging of spiking and subthreshold oscillations in hippocampal neurons**."

| | |
|---|---|
| p-value | 7.82E-04 |
| Wilcoxon rank sum statistic | 121 |

Two-sample Kolmogorov-Smirnov (K-S) tests comparing fluorescence intensity of Archon1 vs. SomArchon at different positions along neurites in cortex, hippocampus, and striatum (see **Supplementary Figure 4**)

**Cortex**

| Distance along neurite from soma ($\mu$m) | p-value |
|---|---|
| 5 | 0.0006 |
| 10 | 0.1451 |
| 15 | 0.0973 |
| 20 | 1.43E-05 |
| 25 | 3.81E-08 |
| 30 | 1.54E-09 |
| 35 | 1.22E-13 |
| 40 | 5.06E-15 |
| 45 | 3.14E-16 |
| 50 | 7.69E-16 |
| 55 | 4.91E-18 |
| 60 | 6.44E-17 |
| 65 | 8.50E-18 |
| 70 | 3.83E-17 |
| 75 | 3.83E-17 |
| 80 | 2.38E-17 |
| 85 | 3.30E-16 |
| 90 | 1.08E-15 |
| 95 | 1.19E-13 |
| 100 | 2.75E-12 |

**Hippocampus**

| Distance along neurite from soma ($\mu$m) | p-value |
|---|---|
| 5 | 0.1543 |
| 10 | 1.54E-05 |
| 15 | 0.0007 |
| 20 | 5.83E-05 |
| 25 | 3.74E-06 |
| 30 | 2.53E-07 |
| 35 | 6.37E-08 |
| 40 | 6.37E-08 |
| 45 | 1.16E-08 |
| 50 | 1.73E-08 |
| 55 | 2.97E-09 |
| 60 | 5.89E-11 |
| 65 | 1.26E-11 |
| 70 | 2.59E-12 |
| 75 | 2.59E-12 |
| 80 | 2.59E-12 |
| 85 | 2.78E-11 |
| 90 | 2.78E-11 |
| 95 | 1.42E-11 |
| 100 | 4.42E-11 |

**Striatum**

| Distance along neurite from soma ($\mu$m) | p-value |
|---|---|
| 5 | 0.1103 |
| 10 | 0.1672 |
| 15 | 0.5598 |
| 20 | 0.1542 |
| 25 | 0.0011 |
| 30 | 8.34E-05 |
| 35 | 5.48E-07 |
| 40 | 5.48E-07 |
| 45 | 1.65E-09 |
| 50 | 5.48E-07 |
| 55 | 1.27E-08 |
| 60 | 1.65E-09 |
| 65 | 1.65E-09 |
| 70 | 8.12E-07 |
| 75 | 1.96E-07 |
| 80 | 3.80E-08 |
| 85 | 6.90E-08 |
| 90 | 1.08E-08 |
| 95 | 1.30E-07 |
| 100 | 4.56E-08 |

Two-sided Wilcoxon Rank Sum test for the membrane resistance of SomArchon-expressing vs. non-expressing neurons in intact mouse brain slices (see **Supplementary Figure 7a**)

| **Membrane Resistance** | **Cortex** |
|---|---|
| p-value | 0.0124 |
| Wilcoxon rank sum statistic | 37 |

| **Membrane Resistance** | **Hippocampus** |
|---|---|
| p-value | 0.6294 |
| Wilcoxon rank sum statistic | 68.5 |

| **Membrane Resistance** | **Striatum** |
|---|---|
| p-value | 0.7308 |
| Wilcoxon rank sum statistic | 52 |

Two-sided Wilcoxon Rank Sum test for the membrane capacitance of SomArchon-expressing vs. non-expressing neurons in intact mouse brain slices (see **Supplementary Figure 7b**)

| **Membrane Capacitance** | **Cortex** |
|---|---|
| p-value | 0.0024 |
| Wilcoxon rank sum statistic | 121 |

| **Membrane Capacitance** | **Hippocampus** |
|---|---|
| p-value | 0.9720 |
| Wilcoxon rank sum statistic | 52 |

| **Membrane Capacitance** | **Striatum** |
|---|---|
| p-value | 0.8357 |
| Wilcoxon rank sum statistic | 51 |

Two-sided Wilcoxon Rank Sum test for the resting potential of SomArchon-expressing vs. non-expressing neurons in intact mouse brain slices (see **Supplementary Figure 7c**)

| **Resting Potential** | **Cortex** |
|---|---|
| p-value | 0.5482 |
| Wilcoxon rank sum statistic | 84.5 |

| **Resting Potential** | **Hippocampus** |
|---|---|
| p-value | 0.7789 |
| Wilcoxon rank sum statistic | 67 |

| **Resting Potential** | **Striatum** |
|---|---|
| p-value | 0.7308 |
| Wilcoxon rank sum statistic | 39 |

Two-sided Wilcoxon Rank Sum test for **Supplementary Figure 8c** between ΔF/F of SomaQuasAr3 vs. SomArchon in intact mouse brain slices

| **ΔF/F** | |
|---|---|
| p-value | 0.0022 |
| Wilcoxon rank sum statistic | 217 |

Two-sided Wilcoxon Rank Sum test for **Supplementary Figure 8d** between SNR of SomaQuasAr3 vs. SomArchon in intact mouse brain slices

| **SNR** | |
|---|---|
| p-value | $1.07\text{x}10^{-4}$ |
| Wilcoxon rank sum statistic | 230 |

**Supplementary Figure 1.** Wide-field fluorescence imaging of live mouse brain slice expressing Archon1-KGC-EGFP-ER2.

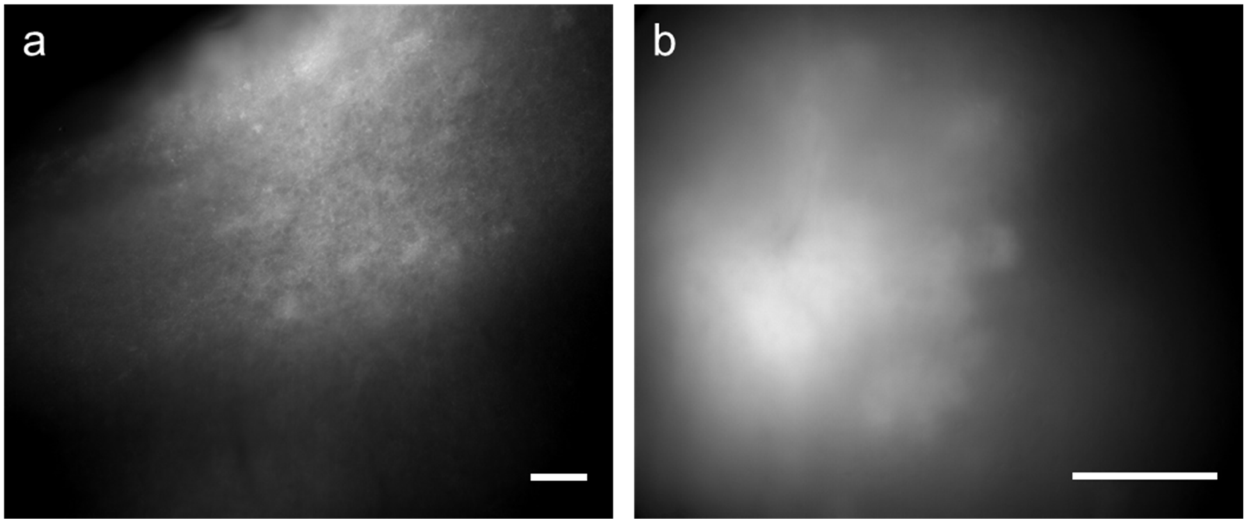


Individual cell bodies of neurons cannot be resolved using high-resolution wide-field microscopy of mouse brain tissue expressing Archon1-KGC-EGFP-ER2 under the CAG promoter, at labeling densities achievable with a standard gene expression technique like *in utero* electroporation (IUE). Fluorescence images of cortical neurons in a 300-µm coronal live brain slice acquired using wide-field microscope with (**a**) 10x and (**b**) 40x objective lens (imaging conditions: excitation 475/34BP from an LED and emission 527/50BP; camera Andor Zyla5.5, binning 1x1; objective lens: 10x NA0.45, 40x NA1.15). Scale bar, 100 µm.

**Supplementary Figure 2.** Population voltage imaging using SomArchon in IUE transduced mouse brain slice.

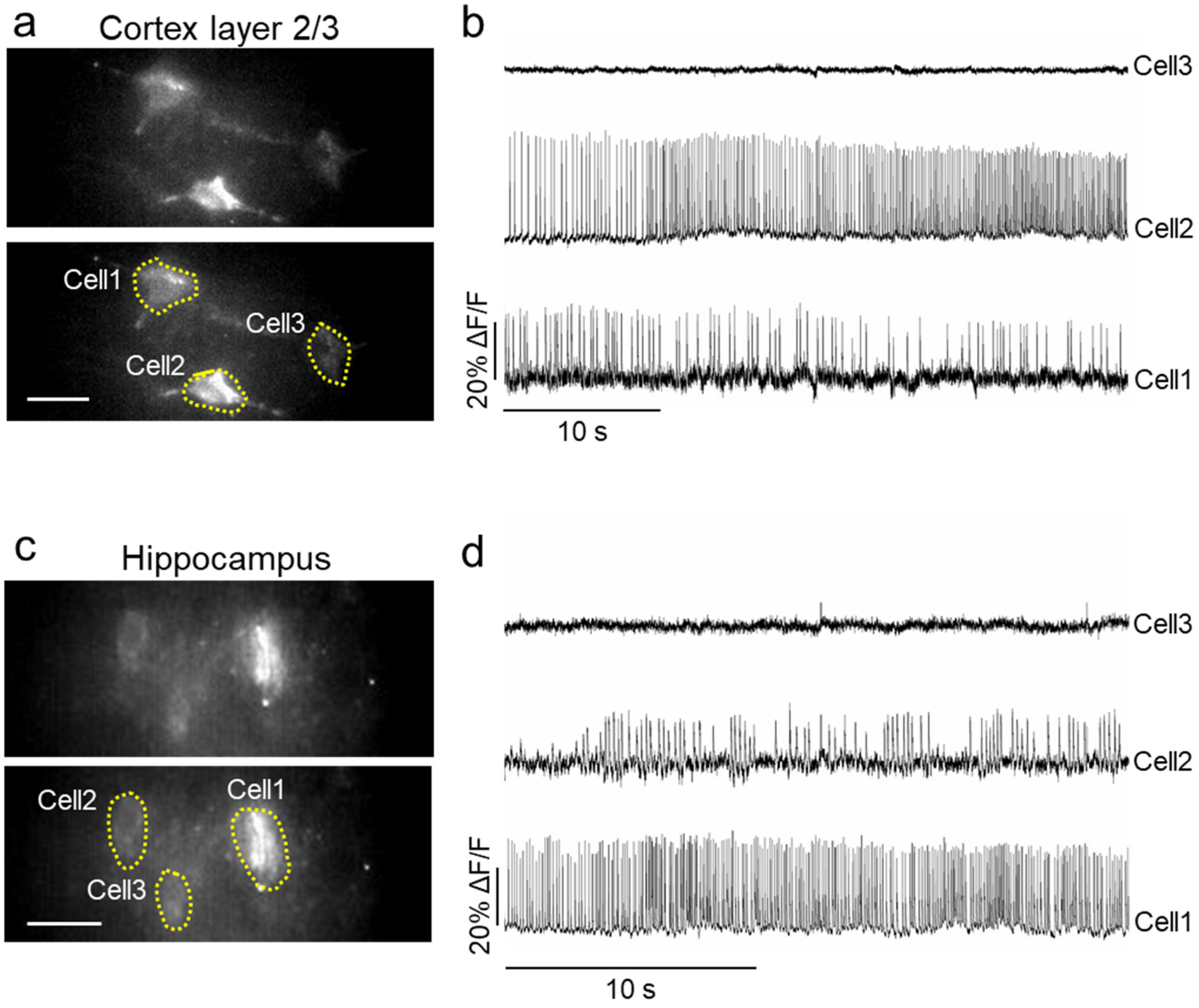


(**a**) Fluorescence wide-field images of a selected field of view in cortex layer 2/3 (top) with selected ROIs corresponding to somata (bottom). Imaging conditions: $\lambda_{ex}$ = 637-nm laser light at 1.5 W/mm$^2$, emission with 664LP, exposure time 2.3 ms. Scale bar, 25 µm. (**b**) Representative single-trial fluorescence traces from cells shown in **a**. Image acquisition rate 440 Hz. (**c**) Fluorescence images of a selected field of view in the hippocampus (top) with selected ROIs corresponding to somata (bottom). Imaging conditions: $\lambda_{ex}$ = 637-nm laser light at 1.5 W/mm$^2$, emission with 664LP, exposure time 3 ms. Scale bar, 25 µm. (**d**) Representative fluorescence traces from cells shown in **c**. Image acquisition rate 333 Hz.

**Supplementary Figure 3.** Population voltage imaging using SomArchon in AAV transduced mouse brain slice.

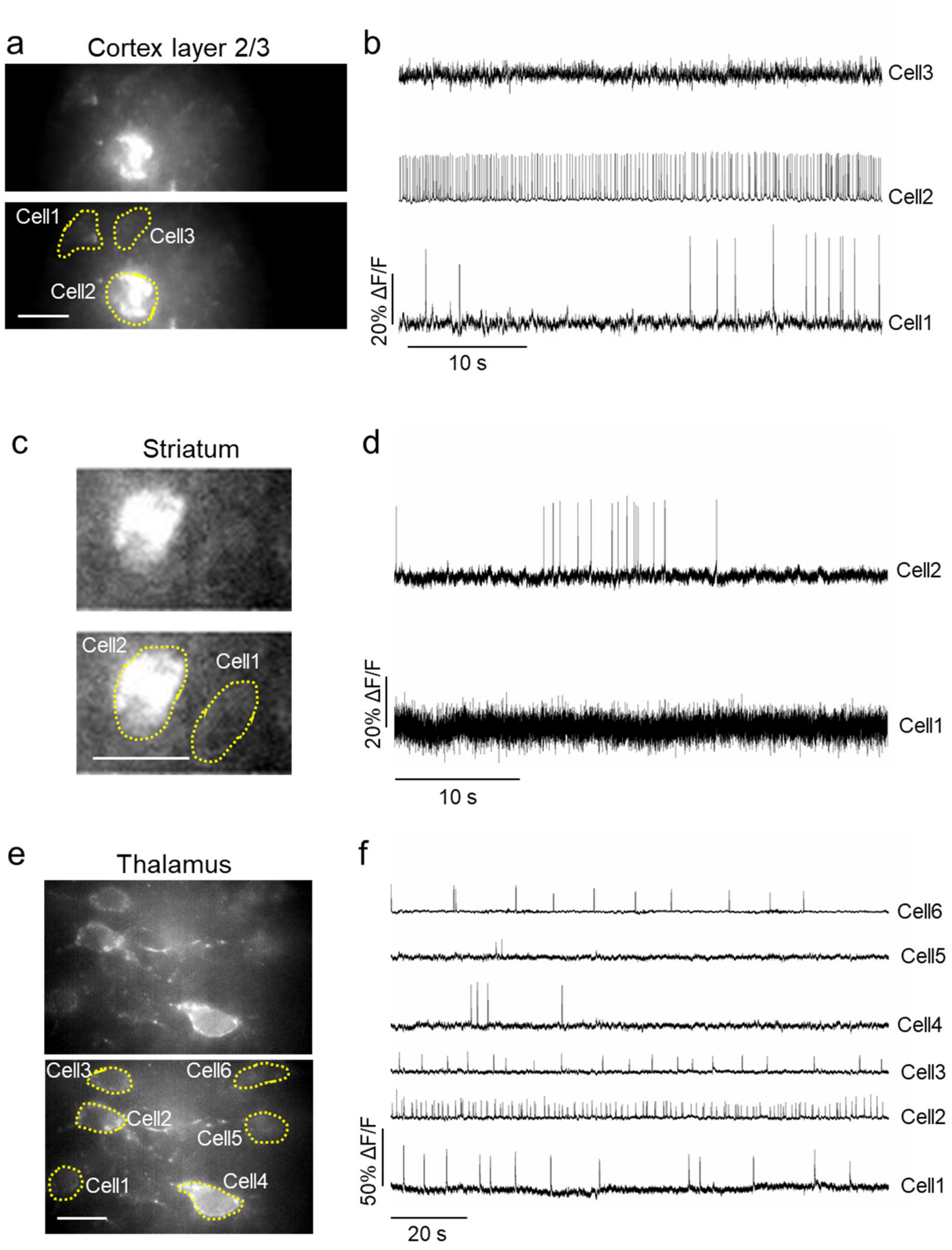


(**a**) Fluorescence wide-field images of a selected field of view in cortex layer 2/3 (top) with selected ROIs corresponding to somata (bottom). Imaging conditions: $\lambda_{ex}$ = 637-nm laser light at 1.5 W/mm$^2$, emission with 664LP, exposure time 1.6 ms. Scale bar = 25 µm. (**b**) Representative single-trial fluorescence traces from cells shown in **a**. Image acquisition rate 632 Hz. (**c**) Fluorescence images of a selected field of view in the striatum (top) with selected ROIs corresponding to somata (bottom). Imaging conditions: $\lambda_{ex}$ = 637-nm laser light at 1.5 W/mm$^2$, emission with 664LP, exposure time 1.4 ms. Scale bar = 25 µm. (**d**) Representative single-trial fluorescence traces from cells shown in **c**. Image acquisition rate 733 Hz. (**e**) Fluorescence images of a selected field of view in the thalamus (top) with selected ROIs corresponding to somata (bottom). Imaging conditions: $\lambda_{ex}$ = 637-nm laser light at 1.5 W/mm$^2$, emission with 664LP, exposure time 3 ms. Scale bar = 25 µm. (**f**) Representative single-trial fluorescence traces from cells shown in **e**. Image acquisition rate 333 Hz.

**Supplementary Figure 4.** Quantification of Archon1 and SomArchon localization in neurons in brain slices.

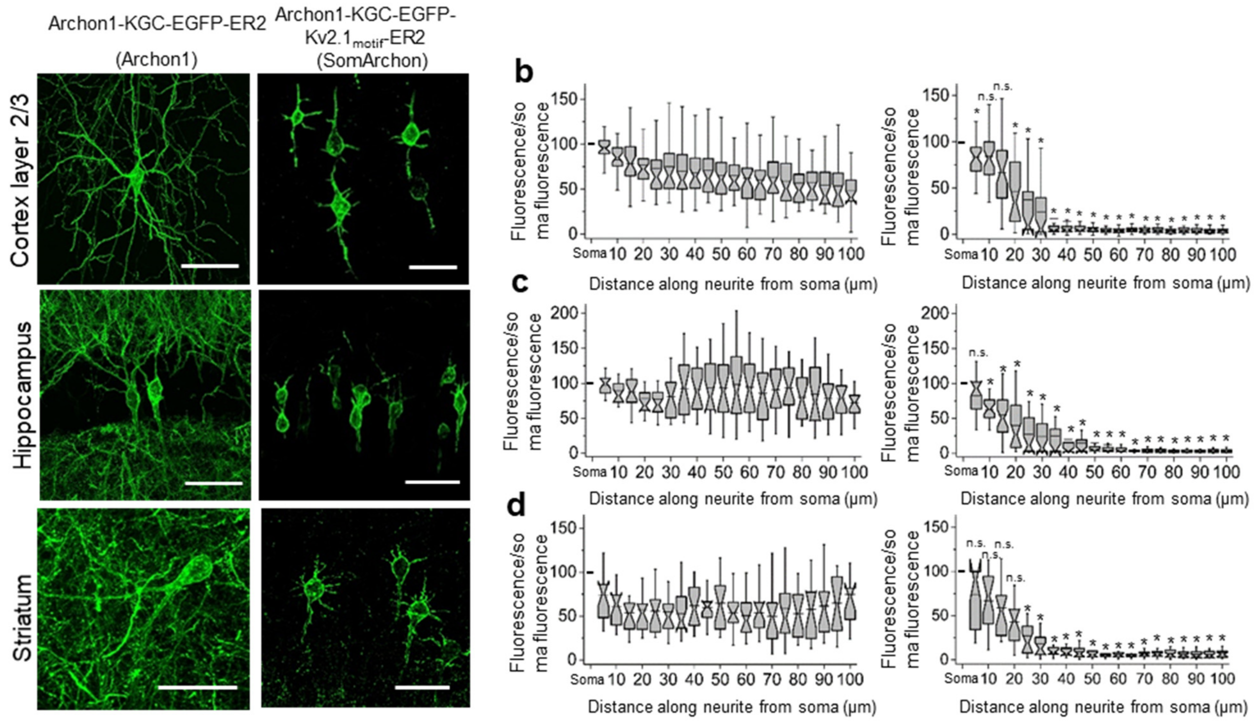


(**a**) Representative confocal images of neurons in cortex layer 2/3 (top), hippocampus (middle), and striatum (bottom) expressing Archon1-KGC-EGFP-ER2 (Archon1; left column) and Archon1-KGC-EGFP-Kv2.1motif-ER2 (SomArchon; right column). Images acquired via EGFP fluorescence using laser excitation at 488 nm and emission 525/50 nm). Scale bars, 50 µm. (**b-d**) Quantification of EGFP brightness versus position along a neurite, normalized to EGFP brightness at the soma, extracted from neurites of Archon1 (left) and SomArchon (right) expressing neurons in (**b**) cortex layer 2/3 (n= 39 and 37 neurites taken from 10 cells from 2 mice each for Archon1 and SomArchon1, respectively), (**c**) hippocampus (n= 20 and 34 neurites taken from 9 and 17 cells from 2 mice each for Archon1 and SomArchon1, respectively), and (**d**) striatum (n= 17 and 20 neurites taken from 7 cells from 2 mice each for Archon1 and SomArchon1, respectively). Box plots with notches are used throughout this paper (narrow part of notch, median; top and bottom of the notch, 95% confidence interval for the median; top and bottom horizontal lines, 25% and 75% percentiles for the data; whiskers extend 1.5× the interquartile range from the 25th and 75th percentiles; horizontal line, mean). P > 0.05 compared to Archon1 at corresponding position away from the soma, not significant (n.s.), throughout all panels of this figure; P* < 0.002 compared to Archon1 at corresponding position away from the soma; two-sample Kolmogorov-Smirnov test. Please see **Supplementary Table 2** for full statistics.

**Supplementary Figure 5.** Expression of Archon1 and SomArchon in cortex and hippocampus.

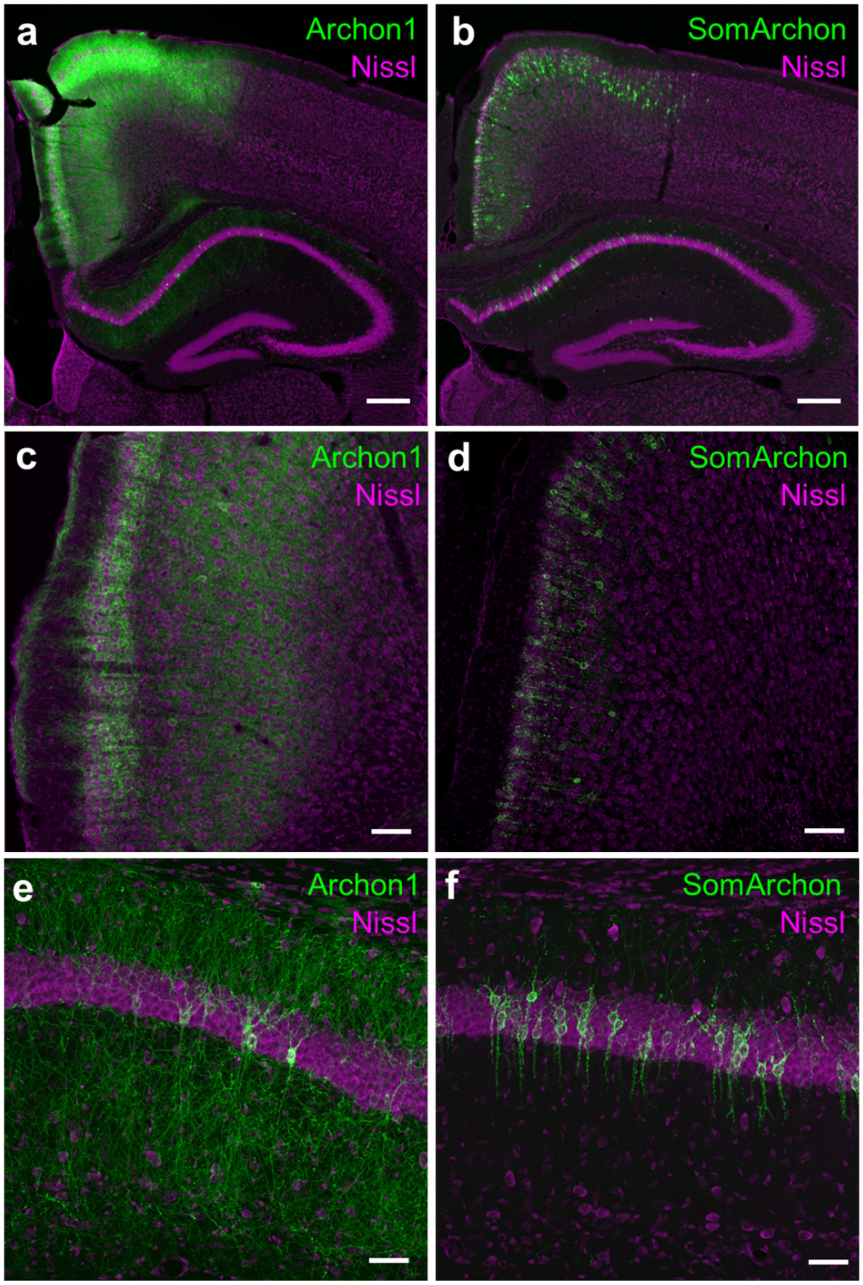


Archon1 or SomArchon expression in mouse brain transduced by IUE at E15.5 and analyzed at P20-P30. (**a-f**) Fluorescence images from 50-μm thick coronal sections of (**a**, **c**, **e**) Archon1- and (**b**, **d**, **f**) SomArchon-expressing brain slices (EGFP channel shown in green; Nissl staining is shown in magenta; near-infrared fluorescence of Archon1 and SomArchon does not survive formaldehyde fixation). (**a**, **b**) Whole brain overview from the hemisphere targeted by IUE demonstrating expression of (**a**) Archon1 and (**b**) SomArchon in neurons in cortex layers 2/3 and hippocampus. Scale bar, 250 μm. (**c-f**) Higher magnification confocal images show expression of (**c, e**) Archon1 and (**d**, **f**) SomArchon in (**c**, **d**) cortex layer 2/3 and (**e**, **f**) hippocampus. Scale bar, 50 μm.

**Supplementary Figure 6.** Expression of Archon1 and SomArchon in striatum.

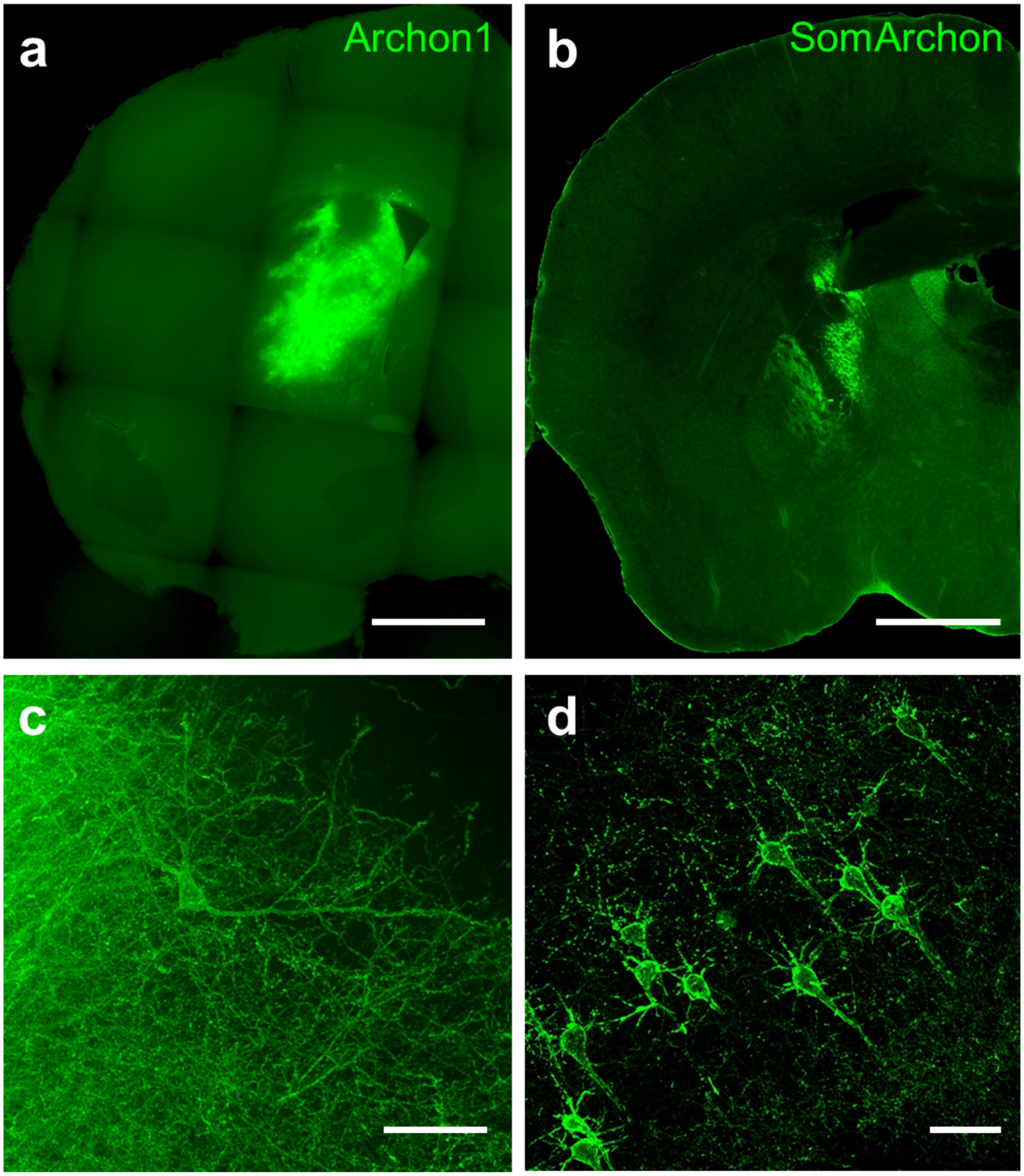


Archon1 or SomArchon were expressed in mouse brain by AAV2-Syn-Archon1 and AAV2-Syn-SomArchon injection. Fluorescence images from 50-μm thick coronal sections of (**a**, **c**) Archon1- and (**b**, **d**) SomArchon-expressing brain slices visualized via EGFP fluorescence (near-infrared fluorescence of Archon1 and SomArchon did not survive formaldehyde fixation). (**a**, **b**) Overview from the hemisphere targeted by AAV injection demonstrating expression of (**a**) Archon1 and (**b**) SomArchon in neurons in striatum. Scale bar, 1 mm. (**c-d**) Higher magnification confocal images show expression of (**c**) Archon1 and (**d**) SomArchon in striatum. Scale bar, 50 μm.

**Supplementary Figure 7.** Membrane properties of neurons in mouse brain slices.

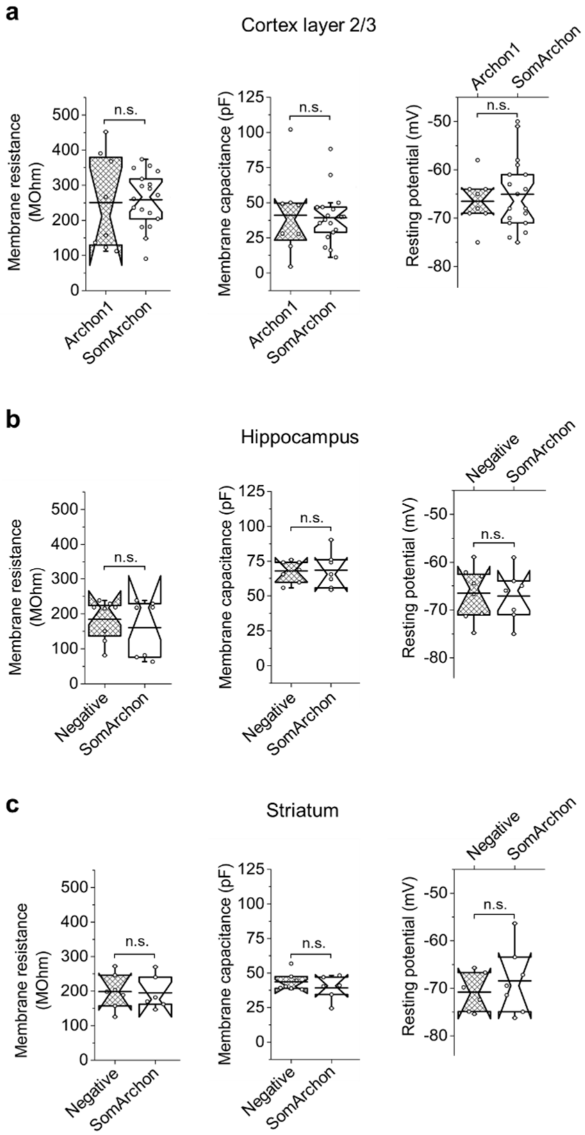


(**a**) Quantification of membrane resistance (left), membrane capacitance (middle), and resting potential (right) of Archon1-expressing (hashed boxes) and SomArchon-expressing (open boxes) neurons in cortex layer 2/3 of live mouse brain slices (n = 8 and 18 cells from 1 and 2 mice for Archon1 and SomArchon, respectively). (**b**) Quantification of membrane resistance (left), membrane capacitance (middle), and resting potential (right) of SomArchon-negative (hashed boxes) and SomArchon-expressing (open boxes) neurons in hippocampus of acute mouse brain slices (n = 8 and 7 cells from 2 mice each for negative and SomArchon, respectively). (**b**) Quantification of membrane resistance (left), membrane capacitance (middle), and resting potential (right) of negative (hashed boxes) and SomArchon-expressing (open boxes) neurons in striatum of acute mouse brain slices (n = 7 and 6 cells from 2 mice each for negative and SomArchon, respectively). Wilcoxon Rank Sum test was performed (see **Supplementary Table 2** for full statistics) to test significant difference.

**Supplementary Figure 8.** Comparison of SomaQuasAr3 and SomArchon in intact brain tissue.

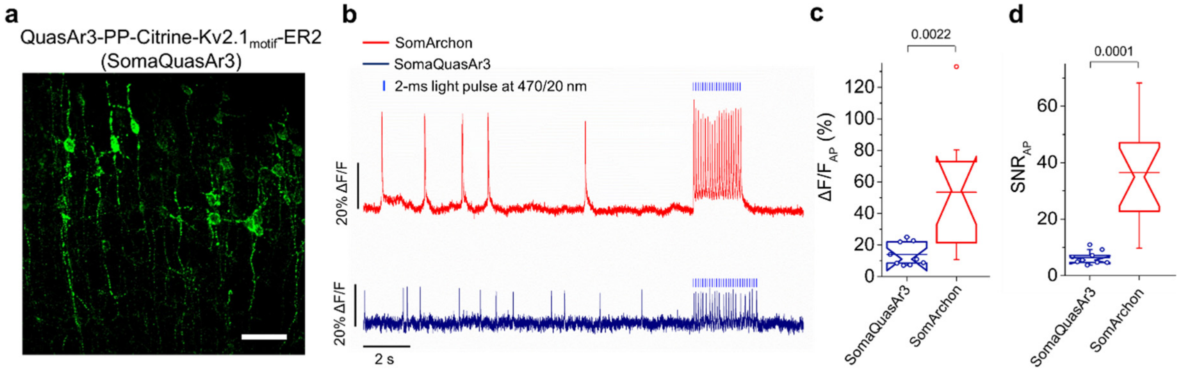


(**a**) Representative confocal image of cortex layer 2/3 neurons expressing QuasAr3-PP-mCitrine-Kv2.1$_{motif}$-ER2 (SomaQuasAr3 for short) visualized via mCitrine fluorescence (laser excitation at 488 nm and emission 525/50 nm). Scale bar, 50 µm. (**b**) Single-trial optical recording of SomArchon (red) and SomaQuasAr3 (dark blue) fluorescence responses co-expressed with CoChR-mTagBFP2-Kv2.2$_{motif}$ during blue light stimulation. Imaging conditions: 637-nm laser, excitation at 1.5 W/mm$^2$, emission with 664LP, acquisition rate ~600 Hz. Blue light (470/20 nm) pulses were 2 ms in duration at 10Hz. (**c**) Quantification of ΔF/F per AP across all recordings performed in cortex layer 2/3 intact brain slices (n = 9 neurons from 2 mice and n = 14 neurons from 2 mice, for SomaQuasAr3 and SomArchon, respectively). p-value from Wilcoxon rank sum test. (**d**) Quantification of SNR per AP across all recordings performed in cortex layer 2/3 intact brain slices (n = 9 neurons from 2 mice and n = 14 neurons from 2 mice, for SomaQuasAr3 and SomArchon, respectively). p-value from Wilcoxon rank sum test.

**Supplementary Figure 9.** Fluorescence images of mouse brain sections prepared after *in vivo* imaging.

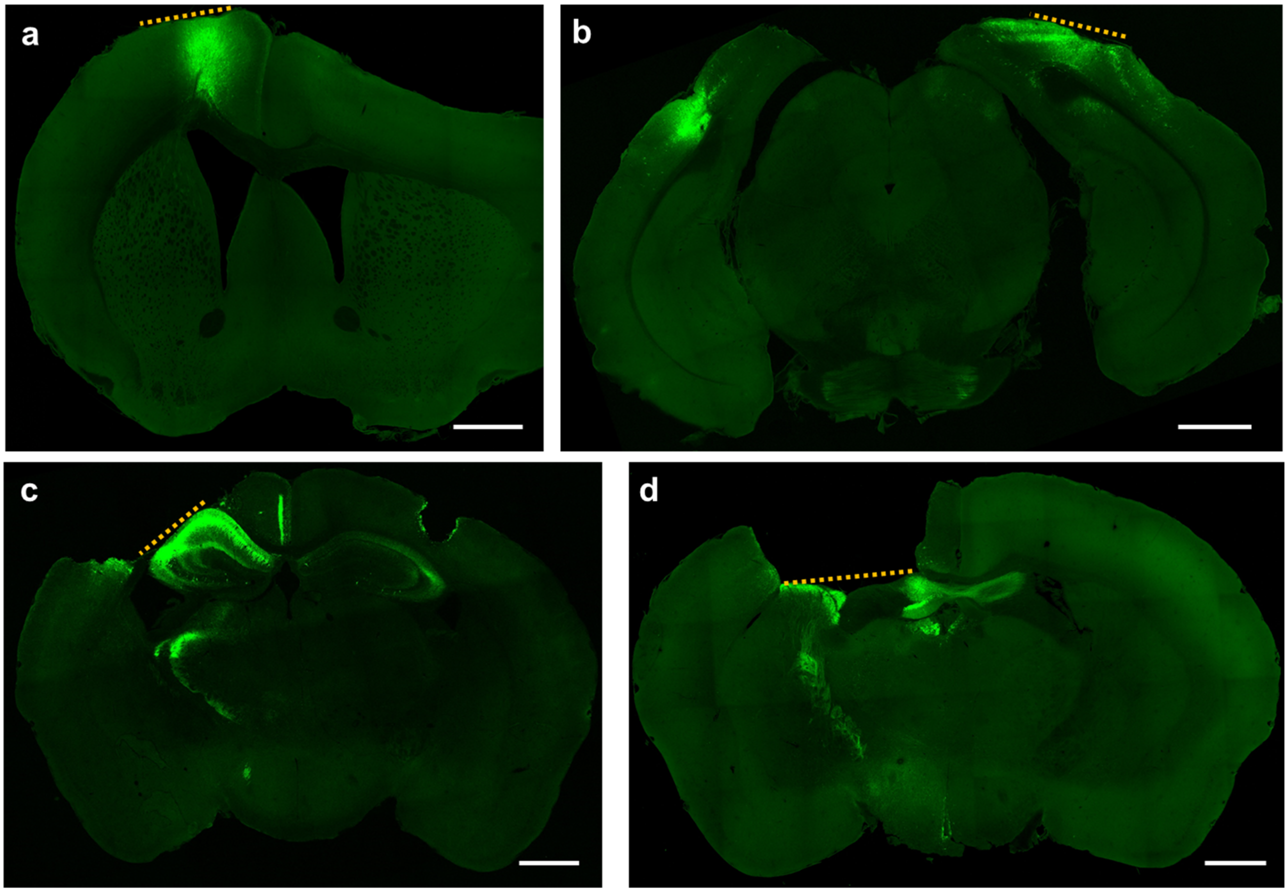


Fluorescence images of brain slices expressing SomArchon in (**a**) motor cortex, (**b**) visual cortex, (**c**) hippocampus, and (**d**) striatum. Dotted line indicates the position of optical imaging window used for *in vivo* imaging. Scale bars, 1 mm.

**Supplementary Figure 10.** Dendritic voltage imaging in the hippocampus of an awake, head-fixed mouse.

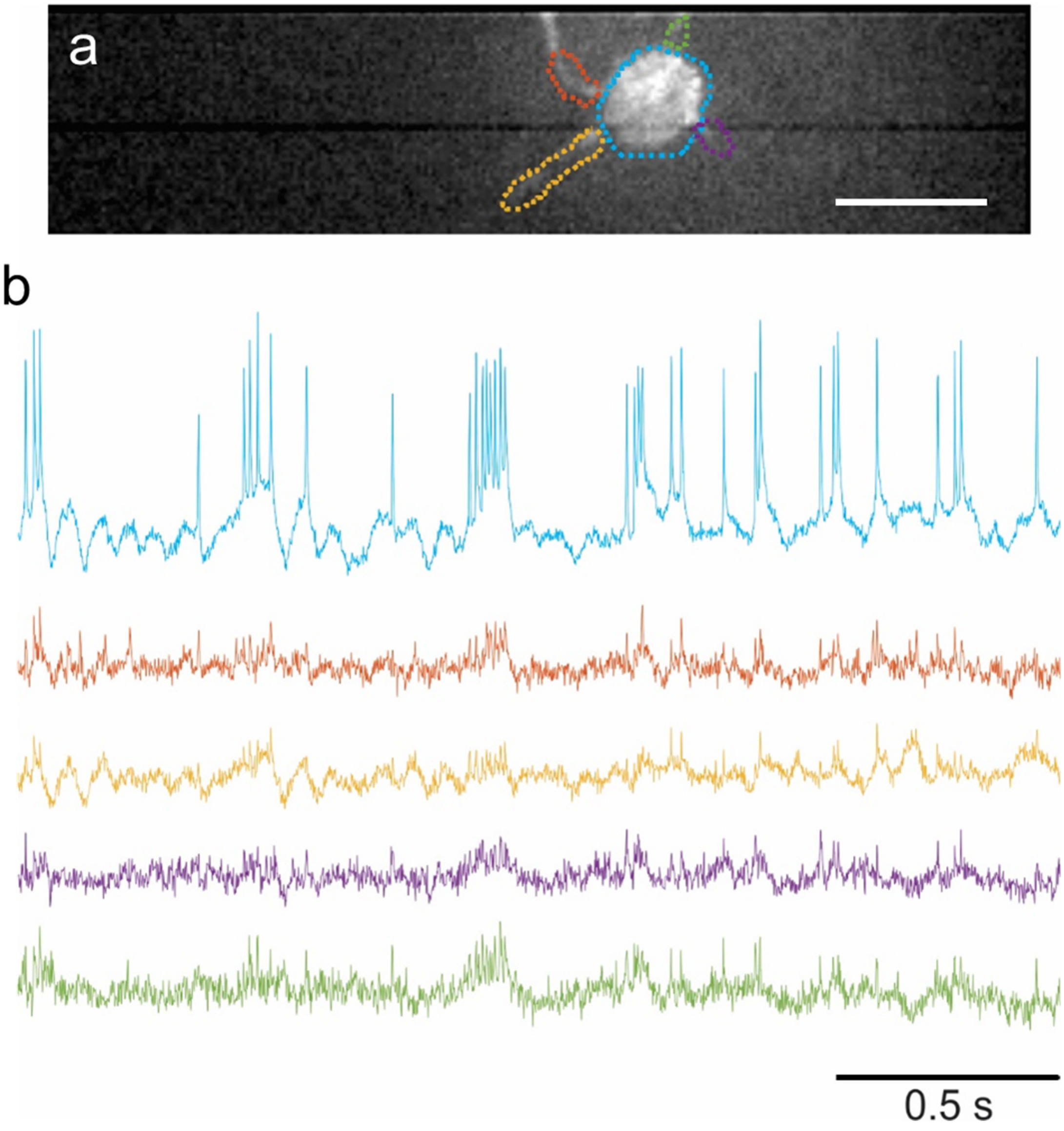


(**a**) Fluorescence image of a hippocampal neuron expressing SomArchon with ROIs selected at the soma and on four proximal dendrites. Scale bar, 25 μm. (**b**) Optical voltage traces from the selected ROIs shown in **a**. Imaging conditions: 637-nm laser excitation at 4 W/mm$^2$; emission 664 nm longpass filter; image acquisition rate: 826 Hz.

**Supplementary Figure 11.** *In vivo* population voltage imaging in primary motor cortex.

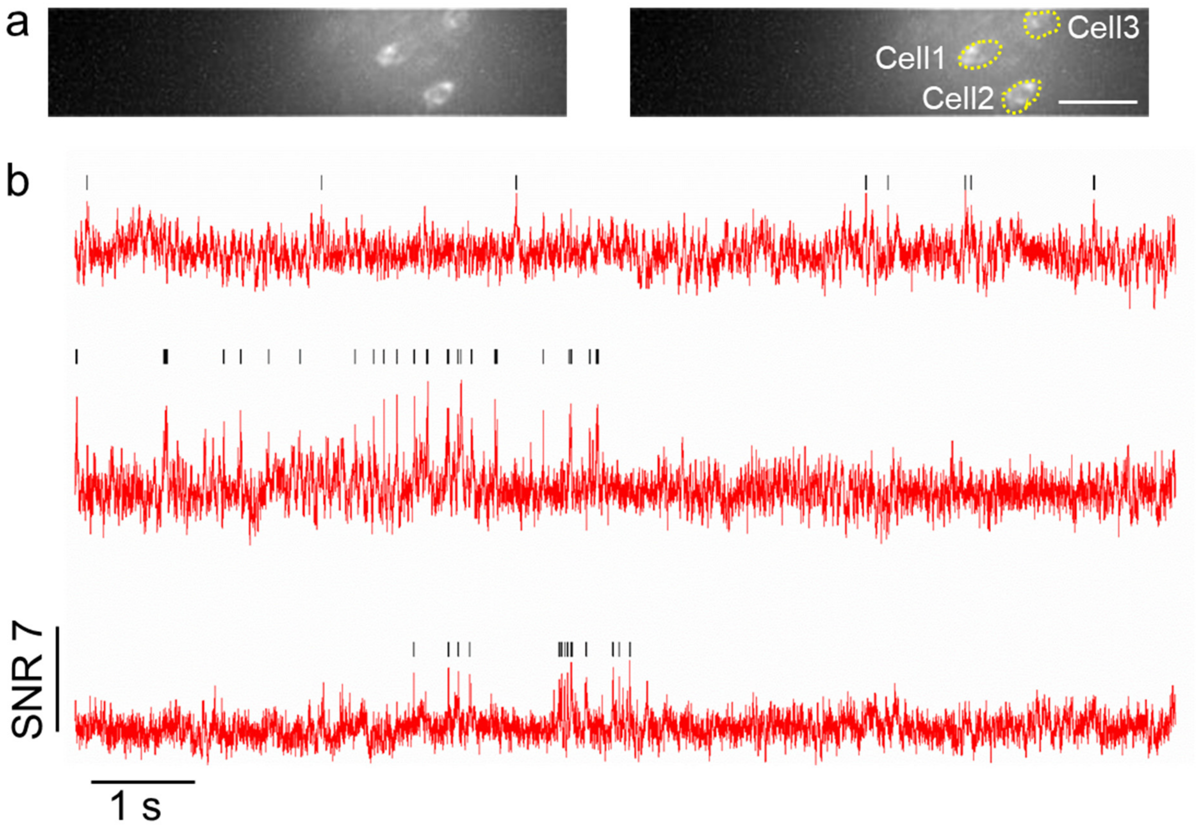


(**a**) Fluorescence images of a selected field of view in motor cortex (left) with selected ROIs corresponding to somata (right), $\lambda_{ex}$=637-nm laser light at 1.5 W/mm$^2$, emission with 664LP, exposure time 1.2 ms. Scale bar = 50 µm. (**b**) Representative fluorescence traces from (**a**) with detected spikes (black line), image acquisition rate 826 Hz.

**Supplementary Figure 12.** Properties of striatal neurons recorded via SomArchon fluorescence.

a

| | Firing Rate (Hz) | Area (um$^2$) | ISI (mean+/- SD) |
|---|---|---|---|
| Cell 1 | 2.00 | 187.59 | 403.8+/-881.5 |
| Cell 2 | 1.6686 | 144.073 | 530.2+/-953.4 |
| Cell 3 | 2.7771 | 172.803 | 300.4+/-542.4 |
| Cell 4 | 2.9167 | 175.338 | 273.3+/-509.6 |
| Cell 5 | 7.933 | 175.76 | 124.4+/-198.3 |
| Cell 6 | 13.2778 | 89.993 | 71.3+/-88.6 |
| Cell 7 | 15.1389 | 69.29 | 64.8+/-87.6 |
| Cell 8 | 11.7222 | 86.612 | 83.7+/-98.9 |
| Cell 9 | 29.95 | 131.82 | 33.2+/-39.8 |
| Cell 10 | 2.0167 | 117.455 | 462.6+/-586.2 |
| Cell 11 | 2.7667 | 149.143 | 351.2+/-470.7 |
| Cell 12 | 5.4271 | 361.66 | 182.4+/-133.1 |
| Cell 13 | 14.3626 | 181.253 | 69.1+/-96.8 |
| Cell 14 | 23.1753 | 191.393 | 43.0+/-44.0 |

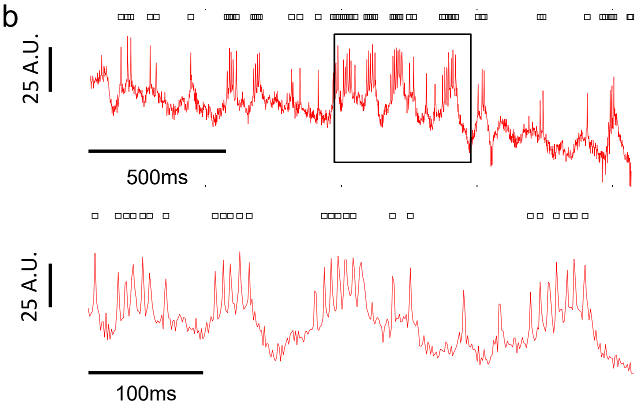


(**a**) Description of the average firing rate, size, and interspike interval (ISI) for the 14 neurons recorded in 9 FOV in 2 mice. Cells recorded in the same FOV are color-coded (blue, red, green). Cells in rows with a white background were recorded individually. We noted that Cells 9 and 14 are bursty, with short ISIs and high firing rates, which are consistent with that of PV cells. Cell 12 is larger in size, with a tonic firing pattern, which is consistent with the characteristics of tonically active cholinergic interneurons. (**b**) Selected trace from Cell 9 exhibiting spike bursting (top). Zoomed view of boxed region in **b** to show spike bursting (bottom). A.U. represents mean fluorescence intensity values.

**Supplementary Figure 13.** Movement speed thresholding for low vs. high speed periods.

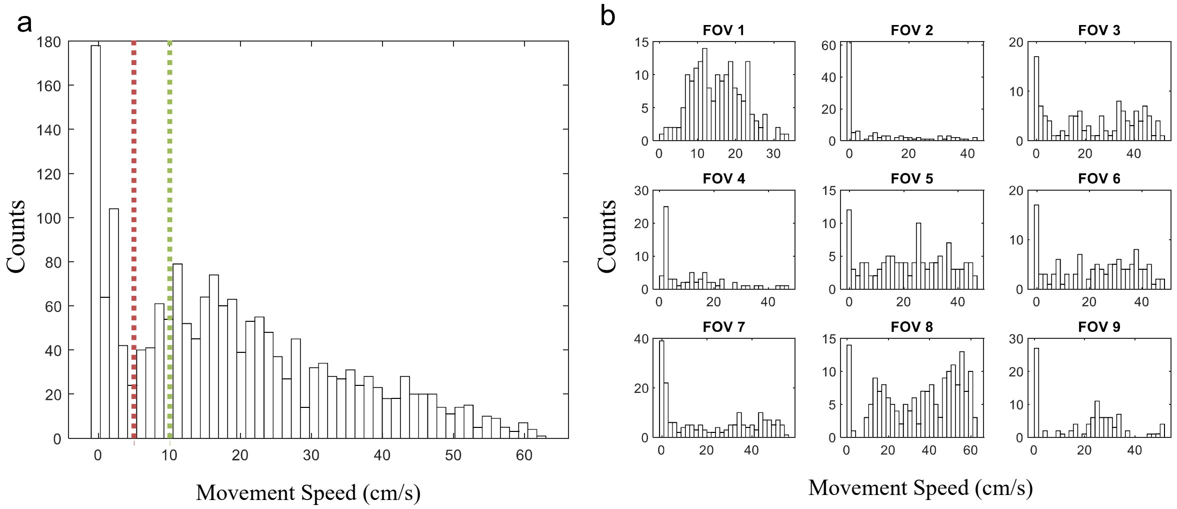


(**a**) Histogram of time intervals with different movement speeds. All recordings were segmented into 0.5-second intervals, and movement speed is calculated as the average speed during every 0.5-second time interval. The red line indicates the threshold for low movement speed identification, and the green line indicates the threshold for high movement speed identification (n=14 neurons in 9 FOVs, n=2 mice). (**b**) Histogram of time intervals with different movement speed for individual FOVs analyzed.

**Supplementary Figure 14.** Eye puff induced changes in LFP high frequency oscillations but not in theta frequency oscillations.

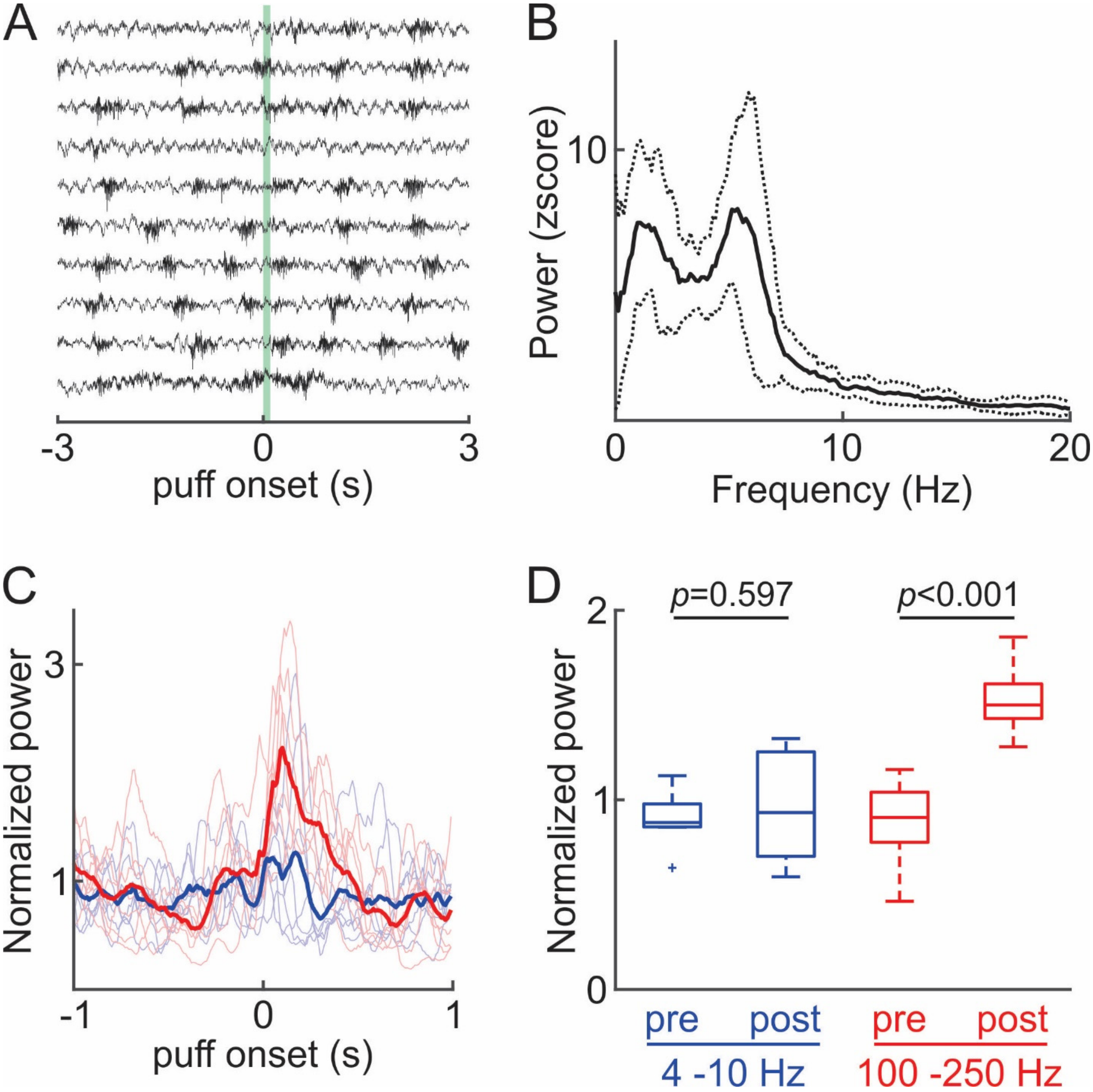


(**a**) An example LFP recording from a session that consisted of 10 trials, aligned to puff onset. Green shading indicates the eye puff. (**b**) LFP power spectrum shows strong theta oscillations (4-10 Hz). (**c**) Changes in oscillation power at high frequencies (100-250 Hz, red) and at theta frequencies (4-10 Hz, blue) induced by puff onset (time 0). Each thin line represents an individual experiment session and the thick lines represent the average across all sessions. (**d**) The eye puff evoked a significant increase in high frequency oscillations (red; $p<0.001$), but not in theta frequency oscillations (blue; $p=0.597$, two-tailed paired Student's t-test, $n=7$ sessions in 4 mice).

**Supplementary Figure 15.** Processing of raw optical voltage traces to remove motion artifacts and global trends.

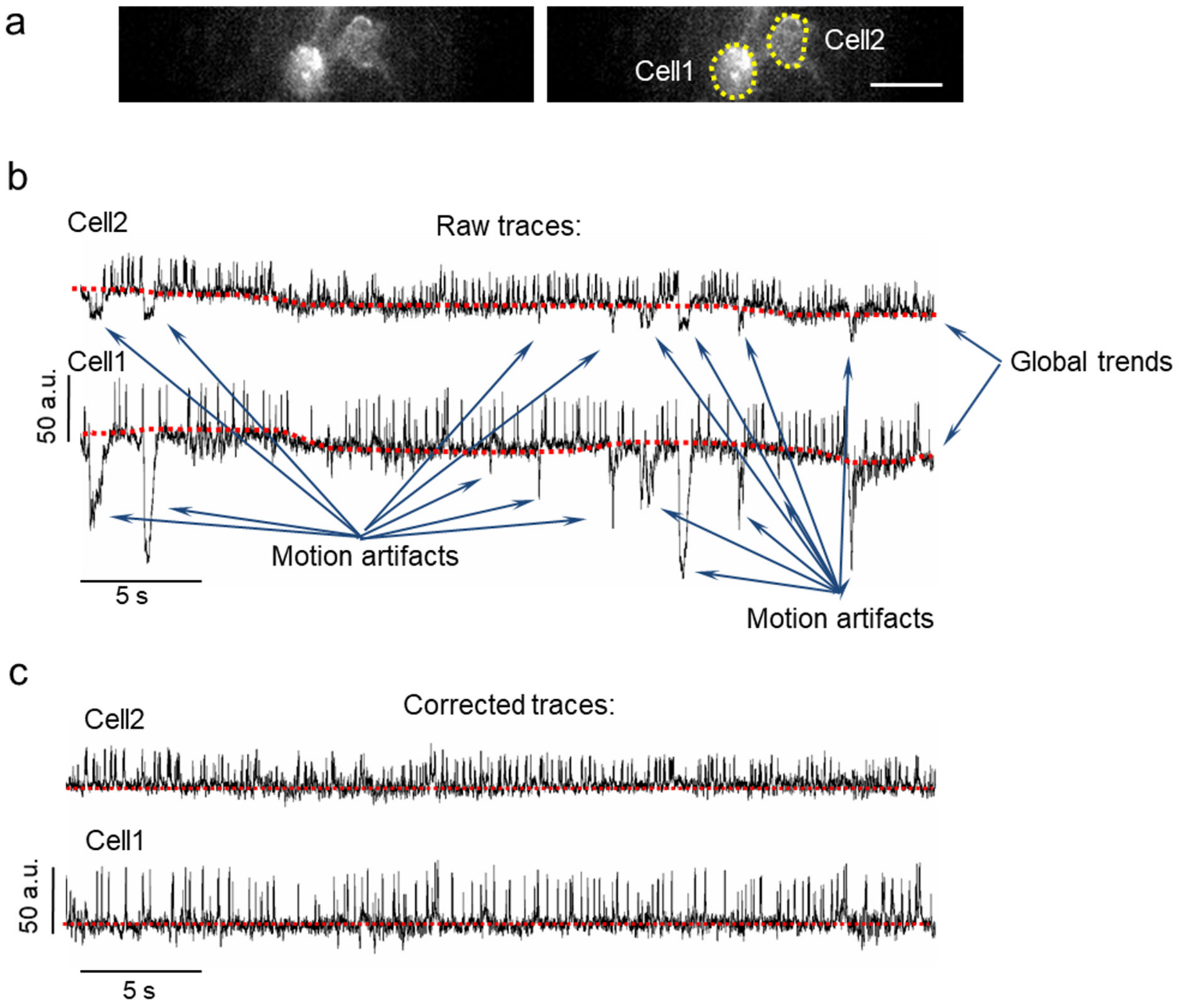


Raw voltage traces were processed before downstream analysis to remove motion artifacts and global trends. Motion results in sharp coordinated changes in fluorescence intensity across all static region of interests for a given field of view. In addition, global trends in baseline fluorescence were observed due to slow photobleaching or slight drift in focus. (**b**) Here, we present raw voltage traces for two cells imaged in the same field of view (as seen in **a**). Both traces demonstrate collateral shifts in fluorescence for both traces as well as common global trend in the baseline shift. (**c**) The motion artifacts in **b** were removed to yield corrected traces.
